# Learning-related population dynamics in right and left dorsal premotor cortex during typing skill acquisition

**DOI:** 10.64898/2026.07.02.736059

**Authors:** Hiroaki Hashimoto, Justin J. Jude, Hadar Levi-Aharoni, Ziv M. Williams, John D. Simeral, Leigh R. Hochberg, Daniel B. Rubin

**Affiliations:** Center for Neurotechnology and Neurorecovery, Department of Neurology, Massachusetts General Hospital, Boston, MA, USA; Department of Neurology, Harvard Medical School, Boston, MA, USA; Department of Neurosurgery, Massachusetts General Hospital, Harvard Medical School, Boston, MA, USA; Harvard-MIT Division of Health Sciences and Technology, Boston, MA, USA; Program in Neuroscience, Harvard Medical School, Boston, MA, USA; School of Engineering, Brown University, Providence, RI, USA; Robert J. and Nancy D. Carney Institute for Brain Science, Brown University, Providence, RI, USA; VA Center for Neurorestoration and Neurotechnology, VA Providence Healthcare System, Providence, RI, USA; Department of Neurological Diagnosis and Restoration, Graduate School of Medicine, The University of Osaka, Suita, Osaka, Japan

## Abstract

Advances in intracortical brain-computer interface (BCI) technology have enabled increasingly sophisticated communication paradigms, including for decoding intended speech and touch typing. However, the methods by which intracortical neural population dynamics are engaged during practice-related skill acquisition in humans remain poorly understood. Here, we examined learning-related changes in neural activity during motor skill acquisition in a right-handed BCI clinical trial participant with tetraplegia, with intracortical microelectrode arrays placed in the bilateral dorsal precentral gyri (Brodmann area 6d), who learned how to type using a BCI-enabled typing interface. While decoder performance remained stable across sessions, typing speed improved with practice, indicating practice-related skill acquisition. Over weeks, low-dimensional neural population activity became progressively more compact, and this compaction was strongly associated with faster typing, independent of decoder accuracy. Although this compaction was observed bilaterally in 6d, firing-rate modulation and cross-session generalization were selectively enhanced in left 6d. Moreover, neural population changes across sessions were largely accounted for by canonical correlation analysis in right 6d, but only partially accounted for in left 6d. Together, these findings demonstrate that human intracortical neuro-motor skill acquisition related to intended typing engages shared bilateral population-level dynamics, with additional learning-related changes selectively expressed in dominant dorsal premotor cortex.

## Introduction

Intracortical brain–computer interfaces (iBCI) have demonstrated the potential to restore reliable communication for individuals with tetraplegia or locked-in syndrome ^1–6^. In the BrainGate clinical trial, participant T18—a right-handed man with cervical spinal cord injury and tetraplegia—recently used an intracortical typing neuroprosthesis that decoded 30 distinct finger movements (2 hands × 5 fingers × 3 gesture types). Using a recurrent neural network (RNN) coupled to a 5-gram language model, the system enabled full bimanual QWERTY keyboard functionality. T18 acquired a typing skill through iBCI and achieved a communication rate of 110 characters per minute (CPM), representing the state-of-the-art neural typing performance ^7^.

In able-bodied individuals, practice of a novel motor task leads to a reduction in movement execution time ^8^. T18 showed a similar improvement, achieving progressively faster neural typing speeds over subsequent recording sessions despite being unable to physically move his fingers. Notably, decoder architecture remained fixed throughout the study, with daily recalibration used solely to compensate for non-stationarities in neural activity. Although the amount of training data increased across sessions, decoding accuracy remained stable, even as T18’s typing speed increased dramatically—from 30–40 CPM at the beginning of training to 110 CPM with practice ^7^. Improvements in typing speed support the acquisition of motor skills. Moreover, as these improvements were not driven by changes in decoder performance, we believe they reflect bona fide learning by the participant.

Previous work has demonstrated that population-level neural dynamics underlying motor learning can be systematically studied using BCI paradigms, primarily in non-human primates. Neural activity during short-term learning is constrained to a manifold, such that learning-related changes occur largely within the intrinsic structure of population activity ^9^. Such manifold-constrained learning can be explained by neural reassociation ^10^, whereby existing patterns of neural activity are remapped to new behavioral outputs without the emergence of fundamentally new population activity patterns, at least when evaluated in a low-dimensional space. In contrast, studies of long-term skill learning have reported the emergence of neural activity patterns that extend beyond the pre-existing manifold ^11^, leading to the conceptual distinction between within-manifold and outside-manifold learning ^12^. In parallel, neural activity associated with an already acquired skill can exhibit day-to-day non-stationarity, yet these changes can be aligned to a manifold across sessions using canonical correlation analysis (CCA) ^13^.

Additional insights into the neural mechanisms supporting rapid motor learning have come from studies examining the interaction between dorsal premotor cortex (PMd) and primary motor cortex (M1), which demonstrate that PMd activity can reorganize within output-null subspaces during rapid learning ^14^. At the single-neuron level, an optogenetic study in mice has further shown that neurons exhibiting strong responses early in learning can decrease their activity as learning progresses, accompanied by increased stability of stimulus-evoked responses ^15^.

Importantly, the majority of these findings have been derived from relatively simple motor tasks, and little is known about how neural population dynamics reorganize during the acquisition of complex motor skills such as typing. In humans, skilled finger-movement learning has been studied primarily using non-invasive methods such as functional MRI ^16–19^, leaving open the question of how population-level principles identified in non-human primates generalize to intracortical recordings in humans. As a result, it remains unclear how long-term skill acquisition in humans reshapes premotor population dynamics, and whether the organizing principles described in animal models extend to complex behaviors enabled by iBCIs.

Here, we analyzed neural population activity recorded from bilateral dorsal premotor cortex during T18’s acquisition of an intracortical typing skill through repeated closed-loop iBCI use. We hypothesized that skill acquisition would be accompanied by systematic changes in premotor neural population dynamics. To test this hypothesis, we examined population activity projected onto task-variable–demixed subspaces using demixed principal component analysis (dPCA) ^20^ and evaluated the extent to which session-to-session changes in neural population activity could be captured within a low-dimensional space using CCA ^13^.

## Results

### Spatiotemporal neural responses to 30 finger movements in right and left 6d

T18 is right-handed, scoring 100% on the Edinburgh Handedness Inventory and 39 on the Waterloo handedness Questionnaire-Revised. Four arrays were implanted in the dominant (left) dorsal premotor cortex (Brodmann 6d) and two arrays in right 6d (Figure 1A), with cortical locations defined here using the Human Connectome Project parcellation ^21^. As a first assessment of neural population tuning, prior to attempting to perform closed-loop typing, the participant attempted to perform 30 distinct finger movements, comprising three gesture types (extend finger up, flex finger down, curl finger into palm) for each finger. Each trial was initiated by the presentation of a written cue followed by a variable preparation period (Figure 1B,C) and a subsequent “Go” signal, which instructed the participant to attempt the cued gesture.

**Figure 1:**
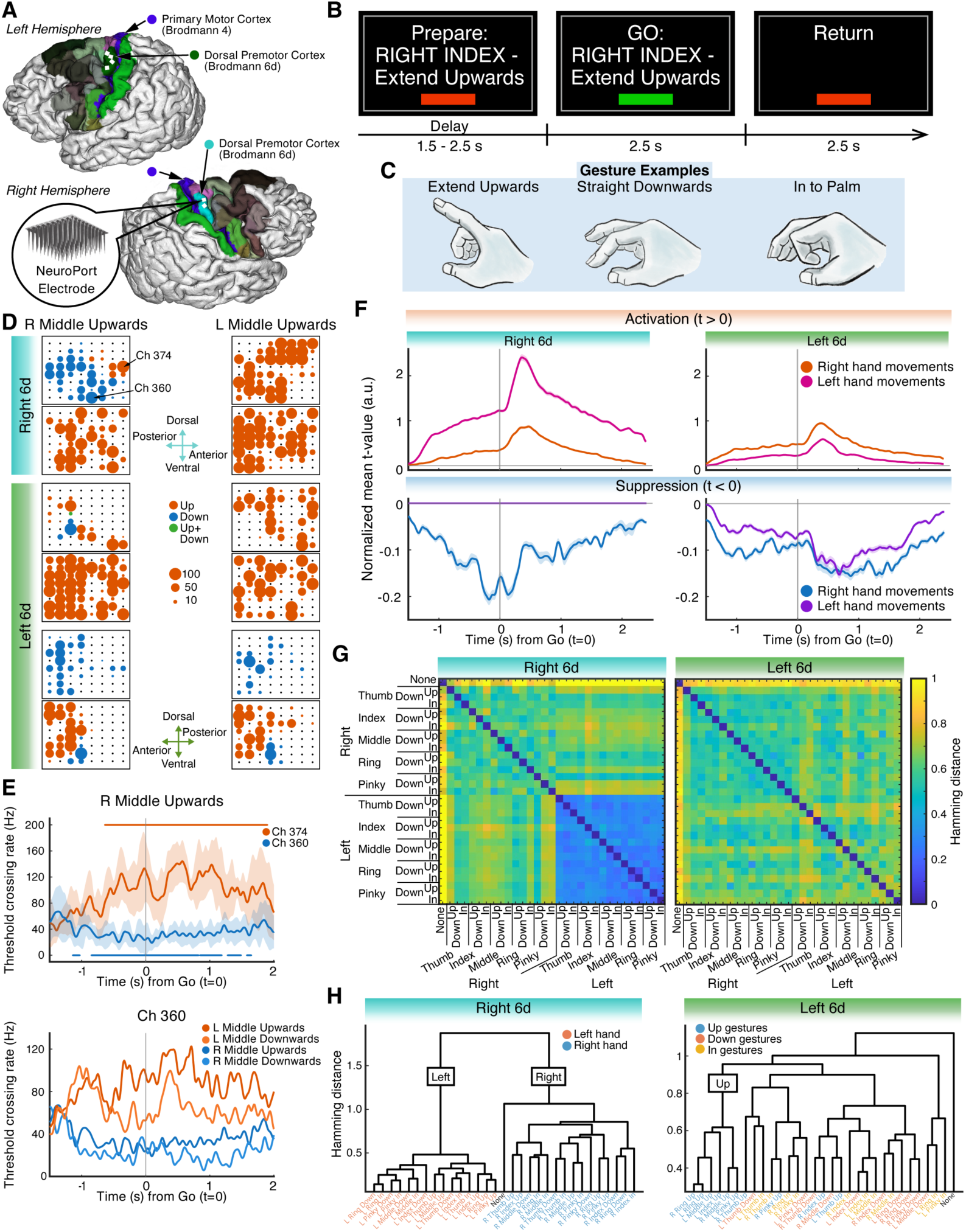
Neural activity in bilateral dorsal premotor cortex (6d) during attempted finger movements. (A) Locations of the implanted NeuroPort electrode arrays in participant T18. Four arrays were placed in the left dorsal premotor cortex (Brodmann 6d, dark green), and two in the right 6d (cyan). The primary motor cortex (Brodmann 4) is shown in blue. White squares indicate electrode arrays. Cortical regions are displayed according to the Human Connectome Project multimodal parcellations. (B) Finger-movement task structure. A variable delay (Prepare) period (1.5–2.5 s) preceded the Go. After the Go, the instructed finger movement was attempted and held for 2.5 s, followed by a Return period instructing T18 to bring the finger back to the resting position. (C) Illustrations of the three instructed finger gestures: extending finger up (Extend Upwards), flexing finger down (Straight Downwards), and curling finger into palm (In to Palm). (D) Spatiotemporal cluster analysis comparing Gaussian-smoothed threshold crossing rates between the Prepare and Go phases. Circles denote channels exhibiting significant increases (red), decreases (blue), or both (green). Circle size reflects the number of significant time points at each channel within 0–1 s after the Go (maximum of 100 time points). (E) Example PSTHs from right 6d during the right middle-finger upward movement. Channel (Ch) 374 shows increased threshold crossing rates, whereas Ch 360 shows decreased threshold crossing rates. Horizontal bars indicate significant time clusters. Bottom: PSTHs of Channel 360 across different finger movements. (F) Normalized mean t-values (from the spatiotemporal cluster analysis) showing activation (*t>* 0) or suppression (*t<* 0) for right-versus left-hand movements in bilateral 6d. Right 6d showed no suppression for left-hand movements. (G) Hamming distance matrices representing the similarity of activity patterns across all attempted finger movements in the right and left 6d. (H) Dendrograms derived from Hamming distances (panel G). All data shown in this figure were obtained during the Finger Movement task on trial day 43.

Before analyzing population structure in reduced-dimensional spaces, we first examined the spatiotemporal organization of Gaussian-smoothed threshold crossing rates derived from non-causal threshold crossings (ncTX) during the 30 finger movements. Throughout the manuscript, these Gaussian-smoothed threshold crossing rates are referred to as firing rate (FR). We computed FR for all 384 channels (64 × 6 arrays) and applied a spatiotemporal cluster analysis ^22^ comparing activity during the Go phase to the Preparation phase. Channels showing significant increases (*t >* 0), decreases (*t <* 0), or both positive and negative responses (*t >* 0 and *t<* 0) were labeled as activation (red), suppression (blue), or mixed (green), respectively (Figure 1D, Supplemental Figure S1). For example, when T18 was instructed to extend his right middle finger upwards, channels on the dorsal array of right 6d exhibited widespread suppression, whereas when instructed to extend his left middle finger upwards, the same region showed broad activation. At the single-channel level, these effects are illustrated by example peristimulus time histograms (PSTHs). Channel 374 displayed activation during right middle finger extend upwards, whereas channel 360 showed suppression during this same movement (Figure 1E, top). Notably, channel 360 exhibited suppression for ipsilateral hand (right-hand) movements but activation for contralateral hand (left-hand) movements (Figure 1E, bottom).

To quantify temporal profiles, channels showing significant activity as identified by the spatiotemporal cluster analysis were selected for each movement, and normalized mean t-values of FR were computed separately for activation and suppression (Figure 1F). Activation showed strong lateralization: in both 6d, contralateral hand movements evoked larger positive normalized mean t-values than ipsilateral movements, consistent with contralateral hand control. Suppression revealed differences between right and left 6d: right 6d showed robust suppression for ipsilateral movements but no suppression for contralateral movements, whereas left 6d showed comparable suppression for both hands.

To examine spatial profiles across movements, each channel was assigned a categorical label (activation, suppression, mixed, or none), yielding a 384-dimensional label vector for each condition. Pairwise masked Hamming distances were computed across all 31 conditions (Figure 1G). In right 6d, movements of the contralateral left hand showed characteristically small distances, indicating that these movements evoked highly similar spatial patterns of activation and suppression. In left 6d, no such clear pattern emerged. Hierarchical clustering demonstrated that right 6d movements separated cleanly by hand, whereas left 6d showed no clear hand-based separation and instead exhibited weak grouping by gesture type (Figure 1H).

### Behavioral and decoding metrics during closed-loop neural typing

Mapping the 30 finger movements to a QWERTY keyboard enabled T18 to perform a closed-loop copy-typing task (Figure 2A). During each trial, the participant attempted to type the prompted sentence, while an online RNN decoded his intended movements in real time. The continuous population activity and the corresponding RNN outputs are shown in Figure 2B (blue line and red shading). The performance of this QWERTY keyboard interface for use as a BCI-enabled communication system has been previously reported ^7^.

**Figure 2:**
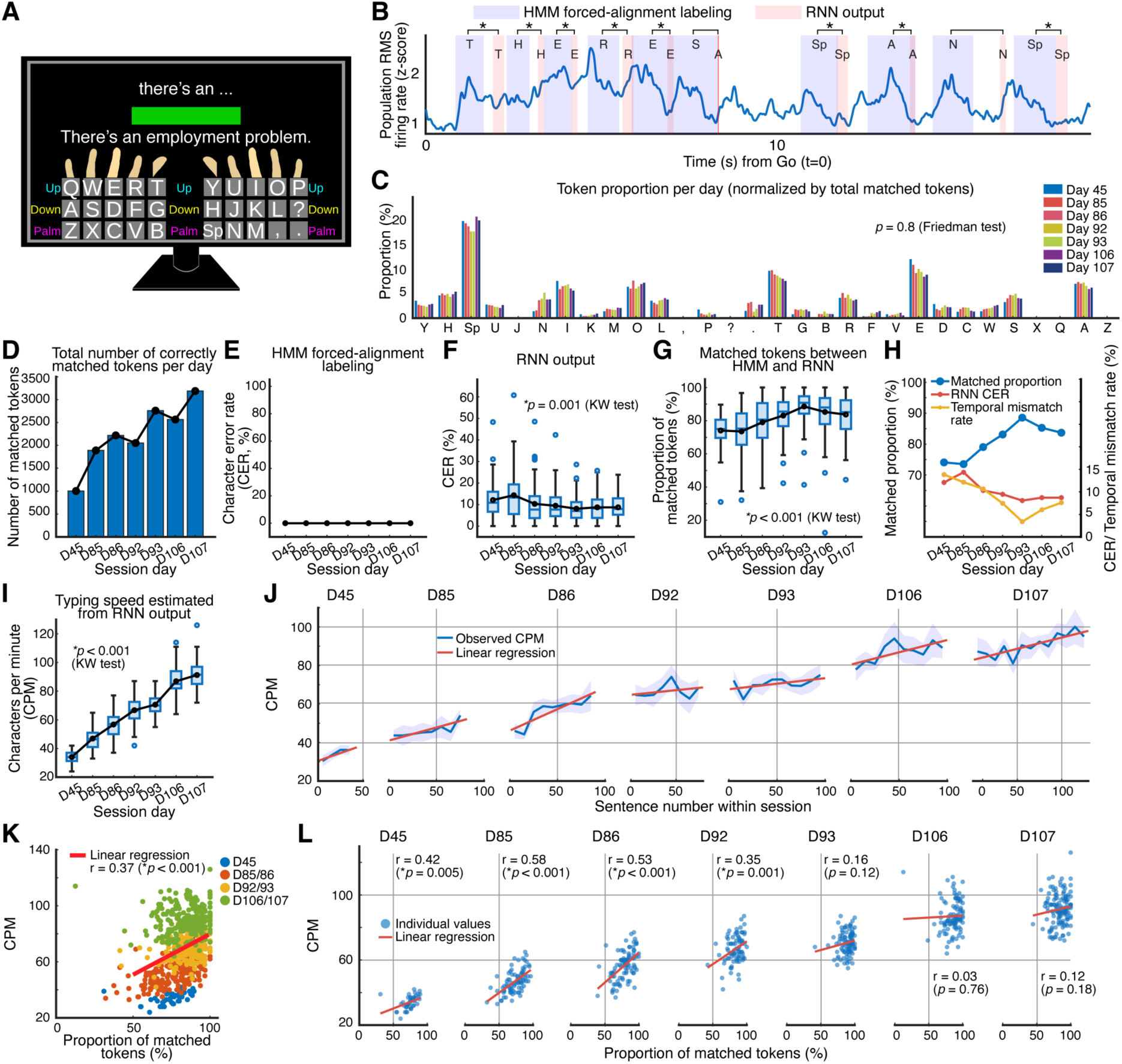
Performance metrics and alignment between HMM and RNN outputs during the closed-loop copy-typing task. (A) Example typing interface. A QWERTY-like keyboard layout was displayed together with the target sentence. Characters were selected via the corresponding finger movements, and intermediate predictions from a 5-gram language model were shown at the top. (B) Example alignment from trial day 45 between HMM forced-alignment labels (blue shading) and RNN output token times (red shading). Tokens were retained (asterisks) when the HMM and RNN token identities matched and their onset times differed by less than 1.5 s. The “N” token was excluded because its HMM and RNN onset times were too far apart (temporal mismatch). (C) Proportion of each token across all closed-loop typing sessions. (D) Total number of tokens correctly matched between HMM labeling and RNN outputs per session. (E) Accuracy of the HMM forced-alignment labeling per session, showing perfect labeling performance (equivalent to a CER of 0%). (F) CER of the RNN output per session. (G) Proportion of matched tokens between HMM and RNN outputs per session. (H) Session-wise temporal mismatch rate computed as: 100% (HMM labeling accuracy) − matched proportion − RNN CER. (I) Typing speed (CPM) estimated from RNN output per session, showing an increase over time. (J) Within-session increase in CPM as a function of sentence number. (K) Positive correlation between typing speed (CPM) and typing accuracy (matched-token proportion) across sessions. (L) Within-session relationship between CPM and matched-token proportion. A significant positive correlation between typing speed and accuracy was present up to Day 92, with reduced correspondence in later sessions.

Because the online RNN decoder does not provide ground-truth timing for individual keystrokes, we applied offline forced-alignment labeling using a Hidden Markov Model (HMM labeling) ^3^ to infer timing of each token (Figure 2B, blue shading). This procedure enabled the extraction of temporally aligned neural activity segments for each token, which were used for subsequent neural analyses (Figure 5, 6, 7) and are described in detail in the Methods. Notably, because the RNN predicts each token using preceding neural activity, its outputs tended to lag behind the HMM labels (Figure 2B). In the present study, the RNN outputs were treated as reflecting the participant’s intended tokens.

We extracted matched tokens—HMM labels that (i) matched the RNN-decoded token identity and (ii) occurred within 1.5 s prior to the RNN output (a hyperparameter chosen to accommodate natural typing variability). Across the seven closed-loop typing sessions, the distribution of token identities remained stable (Friedman test, *p* = 0.8; Figure 2C), while the total number of matched tokens increased monotonically, reflecting the greater total number of sentences presented on later days (Figure 2D).

HMM labeling achieved 100% character accuracy for all sessions (character error rate (CER) = 0%; Figure 2E), reflecting the use of template-based forced alignment with a known, presented character sequence. Although RNN CER differed significantly across sessions (Kruskal–Wallis (KW) test, *p <* 0.001), these differences did not reflect a systematic improvement or degradation over time. Instead, RNN CER values remained within a relatively narrow range across sessions (mean 10.2%, range 8–14%; Figure 2F). The proportion of matched tokens differed significantly across days (KW test, *p<* 0.001) and stabilized at 83–89% during the final sessions (Figure 2G). We defined a temporal mismatch rate—computed as 100% minus the matched-token proportion and the RNN CER—to capture discrepancies between HMM and RNN timings. This rate generally declined across sessions, reaching its lowest value on Day 93 before increasing modestly during the final two sessions (Figure 2H).

Typing speed, measured as the session-wise mean CPM values, differed significantly across sessions (KW test, *p <* 0.001) and increased from 34 to 91 CPM across sessions (Figure 2I), as previously reported for participant T18 ^7^. Typing speed also increased within each session, with CPM rising as the participant progressed through the presented sentences (Figure 2J). We further tested whether more frequently occurring tokens would be typed more rapidly, but found no consistent relationship between token frequency and CPM across sessions (Supplemental Figure S6).

Typing speed was quantified as CPM, while typing accuracy was indexed by the matched-token proportion, defined as the agreement between the keystrokes predicted by the online RNN and the HMM-based ground-truth labels derived from the prompted sentence. Across sessions, CPM correlated positively with matched-token proportion (*r* = 0.37, *p<* 0.001; Figure 2K). Significant within-session correlations were present through D92, but disappeared thereafter as CPM reached a performance ceiling (Figure 2L).

Together, these results show that T18’s improvement in neural typing performance reflected simultaneous gains in both speed and accuracy. This runs counter to the typical speed–accuracy tradeoff and suggests genuine acquisition of motor skill.

### Population-level separability of the 30 finger movements in right and left 6d

Neural activity from the open-loop finger movement task was analyzed to assess the degree to which activity in bilateral 6d was separable among the 30 finger movements. For each channel, class-selective modulation was quantified using false discovery rate (FDR)-corrected KW tests across the 31 classes (30 movements + no movement), applied to three neural features: z-scored ncTX (z-ncTX), z-scored spike-band power (z-PW), and FR. All three features produced substantial numbers of class-selective channels, but z-ncTX and z-PW yielded more KW-significant channels than FR (Figure 3A). Accordingly, subsequent population analyses in Figure 3 used concatenated z-ncTX and z-PW features. Because KW significance only indicates that activity differed across at least one of the 31 classes, these results do not imply that all significant channels exhibited highly selective tuning across finger-movement classes.

**Figure 3:**
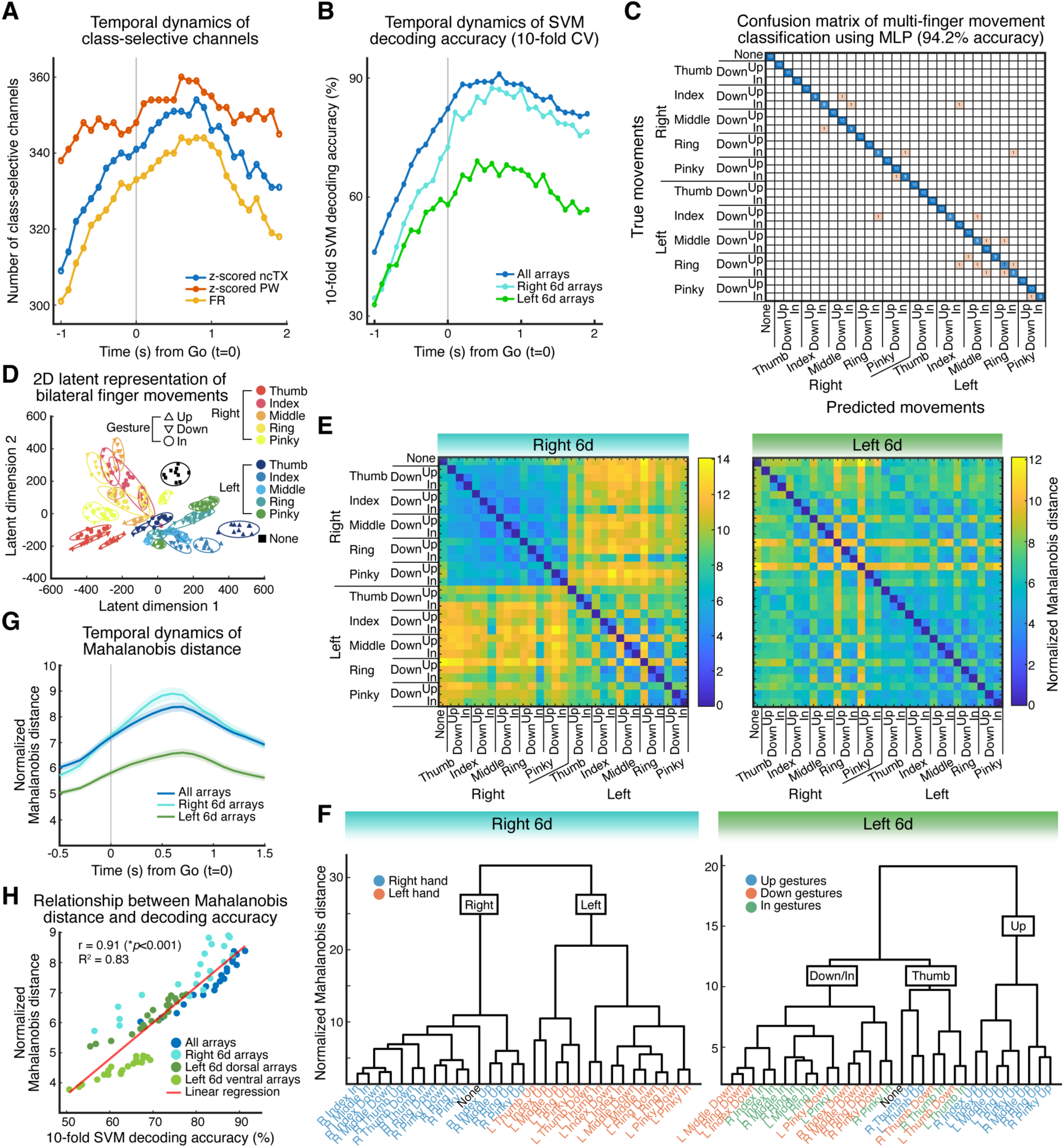
Multivariate neural discriminability and representational structure of bilateral 6d activity. (A) For each channel, neural activity differences across 31 classes (30 finger movements + a no-movement (“none”) class) were assessed using the Kruskal–Wallis test. The time-varying number of channels exhibiting significant differences across classes is shown for each neural feature. (B) Classification accuracy for the 31 classes was computed using an SVM trained on z-scored neural features (ncTX and spike-band power). Decoding performance is shown for all arrays, right 6d, and left 6d across time (10-fold cross-validation). (C) An MLP trained on z-scored neural features (ncTX and spike-band power) achieved a maximum decoding accuracy of 94.2% across the 31 classes. (D) Neural activity patterns for all 31 classes were embedded into a 2D latent space using a trained autoencoder. (E) Pairwise normalized Mahalanobis distances between movement conditions were computed using z-scored neural features (ncTX and spike-band power) and visualized as distance matrices for the right and left 6d. (F) Hierarchical clustering of the distance matrices (panel E) shows that the right 6d primarily encodes hand laterality (right vs. left), whereas the left 6d clusters movements according to gesture type. (G) Time courses of normalized Mahalanobis distances are shown for all arrays, right 6d, and left 6d. (H) Relationship between normalized Mahalanobis distance and SVM accuracy for all arrays, right 6d, and left-6d arrays. The four left-6d arrays are grouped into dorsal (two arrays) and ventral (two arrays) subsets. Each data point corresponds to a time point between −0.5 s and 1.5 s relative to the Go (panel G). All data shown in this figure were obtained during the Finger Movement task on trial day 43.

SVM classifiers were trained using only KW-significant channels to discriminate the 31 classes, and classification accuracy was highest when using all six arrays (Figure 3B). A multilayer perceptron (MLP) trained on all-array data achieved 94.2% accuracy (Figure 3C). We then used this MLP architecture as the basis for an autoencoder to project neural activity into a two-dimensional latent space ^23^. In this latent space, the “none” condition formed an isolated cluster, whereas movements of the right and left hands occupied opposing regions along a diagonal axis of the manifold (Figure 3D).

To quantify representational geometry directly from neural features, we computed pairwise normalized Mahalanobis distances between all condition pairs separately for right and left 6d (Figure 3E). Hierarchical clustering of these distances revealed a clear hand-based structure in right 6d, whereas left 6d was organized primarily by gesture type (Figure 3F). Notably, this organization closely mirrored that observed in Figure 1H based on spatial activation patterns.

Across time, normalized Mahalanobis distances peaked shortly after the Go (Figure 3G). Using time-resolved values computed at each pre- and post-Go time point, the mean normalized Mahalanobis distance showed a significant positive correlation with SVM classification accuracy (*r* = 0.91, *p <* 0.001; Figure 3H), indicating that larger representational separation supports more accurate decoding.

### Distinct low-dimensional population dynamics in right and left 6d

Neural activity during open-loop 30 finger movements was projected into a low-dimensional space using two complementary dimensionality-reduction methods—jPCA ^24–26^ and dPCA ^20,27^ —applied to z-scored FR (z-FR).

Using jPCA, we observed significant rotational dynamics shared across all finger movements, with a brief peak using a 200-ms window centered 0.25 s after the Go (Supplemental Figure S4). We next used dPCA to dissociate contributions of five marginalizations—Hand, Finger, Gesture, Common(i.e., Time), and their interactions. Right 6d primarily encoded Hand information (Figure 4A, C), whereas left 6d encoded mainly Finger and Gesture signals (Figure 4B, D), consistent with the spatial organization observed in Figure 1H and Figure 3F. The dPCA analysis was repeated separately for each implanted electrode array, and consistent patterns were observed across individual arrays within each right or left 6d (Supplemental Figure S7).

**Figure 4:**
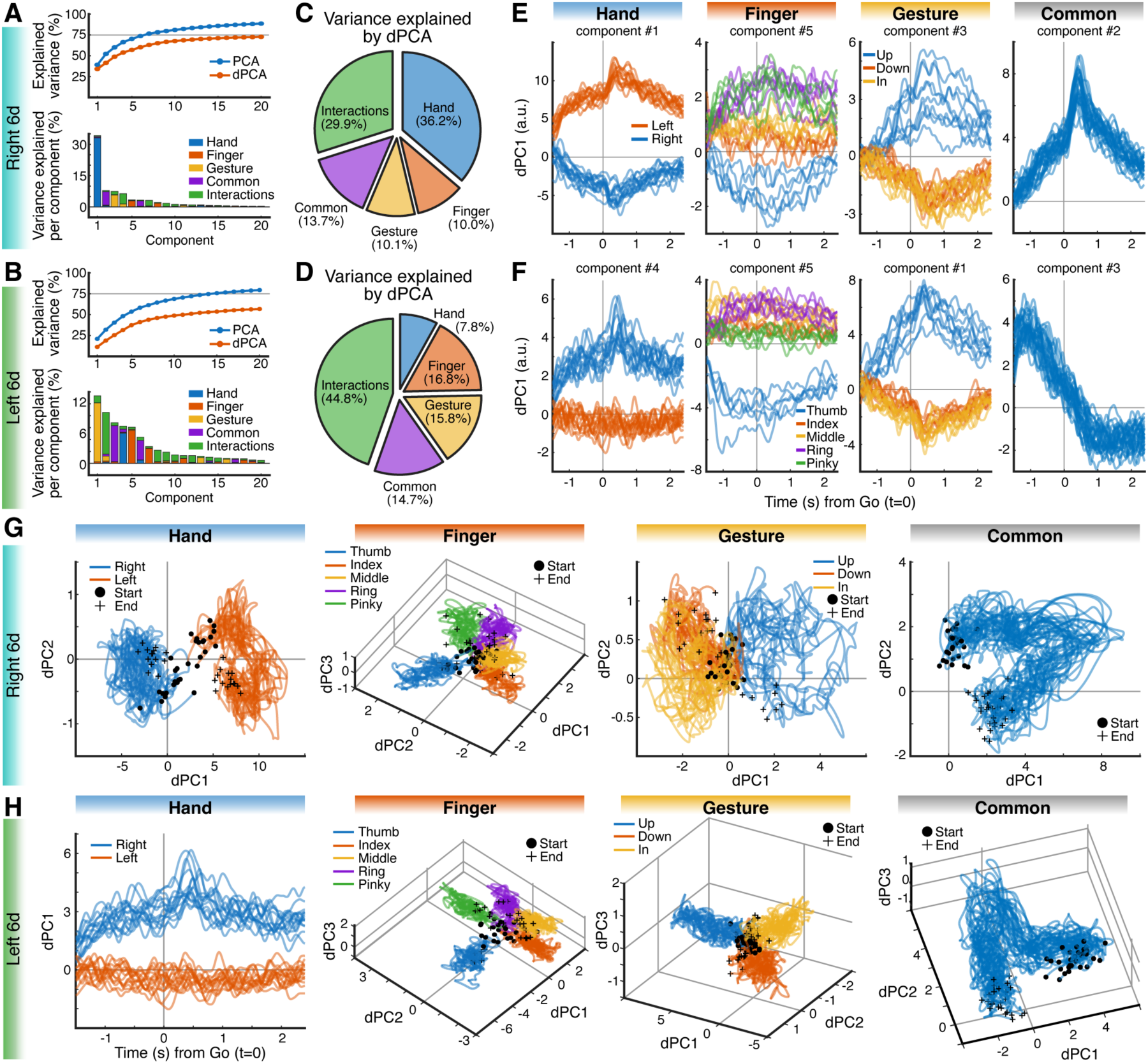
Low-dimensional demixed neural representations of finger movements in bilateral 6d. (A, B) Cumulative variance explained by PCA (blue) and dPCA (red) for right 6d (A) and left 6d (B) (top). Fraction of variance per component attributed to Hand, Finger, Gesture, Common, and Interactions marginalizations (bottom). Analyses were performed using z-scored FR across all 30 finger movements. (C, D) Overall variance distribution across marginalizations obtained from dPCA for right 6d (C) and left 6d (D). (E, F) Time courses (−1.5 to 2.4 s from the Go) of the top demixed components for each marginalization in right 6d (E) and left 6d (F). Hand-related components show contralateral-dominant activity, with positive responses for the controlled (opposite) hand. (G, H) Neural trajectories projected onto the top demixed components (1–3) for right 6d (G) and left 6d (H). Trajectories begin at −1.5 s (black dots) and end at 2.4 s (crosses). For left 6d, the Hand marginalization produced only one dPC; thus, the Hand trajectory is identical to the time course shown in panel F. All data shown in this figure were obtained during the Finger Movement task on trial day 43.

The temporal profiles of the top component for each marginalization are shown in Figure 4E, F. Notably, Hand components exhibited clear lateralization: both 6d showed positive values for contralateral movements and negative values for ipsilateral movements.

Because each dPCA component reflects a mixture of marginalizations, we assigned each component to the marginalization that explained the largest fraction of its variance. Trajectories were plotted along the extracted components (range of 1 to 3 components) for each marginalization (Figure 4G, H). In both 6d, the arrangement of finger trajectories in latent space mirrored the anatomical organization of the digits. Gesture structure indicated a two-dimensional representation in right 6d and a three-dimensional representation in left 6d, with the three gestures forming well-separated clusters in left 6d.

Together, these analyses show that right 6d was dominated by Hand information, whereas left 6d emphasized Finger and Gesture representations. Thus, the bilateral 6d exhibited distinct representational structure rather than mirror-symmetric representations.

### Token-aligned neural activity reveals preserved movement structure and laterality-dependent changes in 6d

To validate the token-aligned neural activity, we segmented z-FR using HMM-labeled token timings. Because the raw segments varied in duration, token-wise FR was temporally aligned and refined using a time-warping procedure (Supplemental Figure S8) ^28^.

Representational structure was quantified using normalized Mahalanobis distances across token classes, computed separately for right and left 6d within each session (Figure 5A). Right 6d dendrograms demonstrated a consistent hand-based organization across days (Figure 5B, Supplemental Figure S9), indicating that the HMM-based segmentation produced stable and behaviorally meaningful neural patterns.

**Figure 5:**
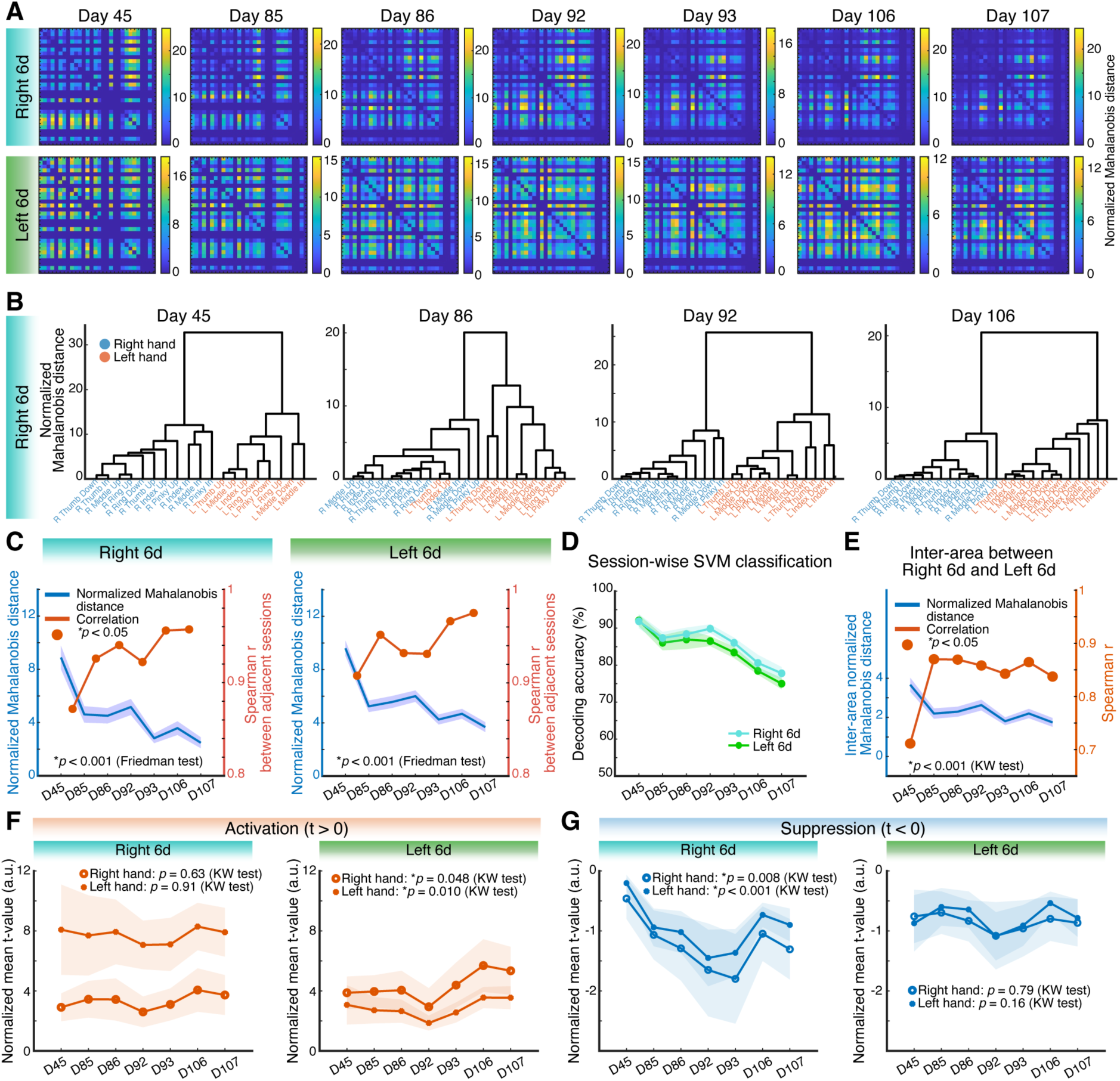
Session-by-session analysis of neural representations derived from HMM-segmented finger-typing activity in bilateral 6d. (A) Session-wise normalized Mahalanobis distance matrices computed from HMM-segmented neural activity (z-scored FR) for 30 finger-movement tokens. Unused token positions were filled with zeros. Labels are omitted for clarity but correspond to the ordering used in Figure 3E, excluding “None.” Matrices are shown for right 6d (top) and left 6d (bottom) across all closed-loop typing sessions. (B) Hierarchical clustering of normalized Mahalanobis distances for right 6d on selected days (Day 45, 86, 92, and 106). Across sessions, right 6d consistently clusters finger movements according to hand laterality (right vs. left hand). (C) Session-wise changes in the mean normalized Mahalanobis distance (blue; 95% CI) for right and left 6d. Both regions showed a decreasing pattern across sessions, with the Friedman test indicating significant differences across days (*p <* 0.001). Spearman correlations of normalized Mahalanobis distances between adjacent sessions (red) show increasing inter-session similarity, with significant correlations marked by red circles (*p<* 0.05). (D) Session-wise SVM decoding accuracy for classifying the finger movements using 10-fold cross-validation. Both right and left 6d exhibit decreasing decoding accuracy over days. (E) Inter-area comparison of representational structure between right and left 6d. The absolute difference in normalized Mahalanobis distances (blue; 95% CI) shows a marked decrease from Day 45 to Day 85, during which the Spearman correlation between areas (red) increases substantially, with significant correlations indicated (*p<* 0.05). (F) Normalized mean t-values for activation (*t>* 0), computed by multiplying the mean t-value by the proportion of significantly modulated channels from spatiotemporal cluster analysis. Right 6d shows no significant session-wise change. In left 6d, activation-related values increased for both right-hand and left-hand movements, with significant session-wise differences detected by the Kruskal–Wallis test (*p<* 0.001). (G) Normalized mean t-values for suppression (*t<* 0). Left 6d exhibits no significant changes across sessions, whereas right 6d shows decreased suppression-related values for both right-hand and left-hand movements, with significant session-wise differences detected by the Kruskal–Wallis test (*p<* 0.001).

Across sessions, mean normalized Mahalanobis distances decreased in both 6d, and correlations between Mahalanobis distance matrices from adjacent days increased (Figure 5C), indicating a progressive compaction of the internal representational geometry within each area. SVM classification accuracy was computed offline within each session using cross-validation, independent of the online RNN decoder. Consistent with the positive relationship between normalized Mahalanobis distance and SVM accuracy (Figure 3H), decoding accuracy declined over days as normalized Mahalanobis distances decreased (Figure 5D). However, despite this decline, classification performance remained far above chance level (3.3% for 30 classes), indicating that neural population activity continued to exhibit substantial separability across tokens. The inter-area difference in representational structure, quantified as the absolute difference in mean normalized Mahalanobis distance between right and left 6d, also decreased across days; notably, right and left 6d became more correlated between Day 45 and Day 85 (Figure 5E).

Neural activity during typing versus rest was compared using spatiotemporal cluster analysis (Supplemental Figure S10), calculating normalized mean t-values for activation (*t >* 0) and suppression (*t <* 0). Activation exhibited robust lateralization: in both 6d, contralateral-hand movements showed consistently higher activation than ipsilateral hand movements (Figure 5F) —matching the pattern observed in Figure 1F. However, in the dominant left 6d, activation values differed significantly across sessions (KW tests), and inspection of the session means revealed an overall increase from the initial to the final day, whereas right 6d remained stable. Conversely, suppression showed significant changes in right 6d but not left 6d, with stronger suppression emerging on later days (Figure 5G). Importantly, population geometries became increasingly similar across sessions and across premotor areas, whereas right and left exhibited distinct FR–based adaptations.

### Connectivity analysis across sessions

Token-aligned PSTHs were computed from HMM-segmented z-FR, and principal component analysis (PCA) was subsequently performed. To separate signal from noise, we generated a surrogate distribution and identified significant principal components (PCs) through noise-corrected dimensionality estimation ^14^. Both 6d yielded a median of eight signal PCs for right-hand movements, whereas for left-hand movements, the contralateral right 6d showed significantly higher dimensionality (Figure 6A).

**Figure 6:**
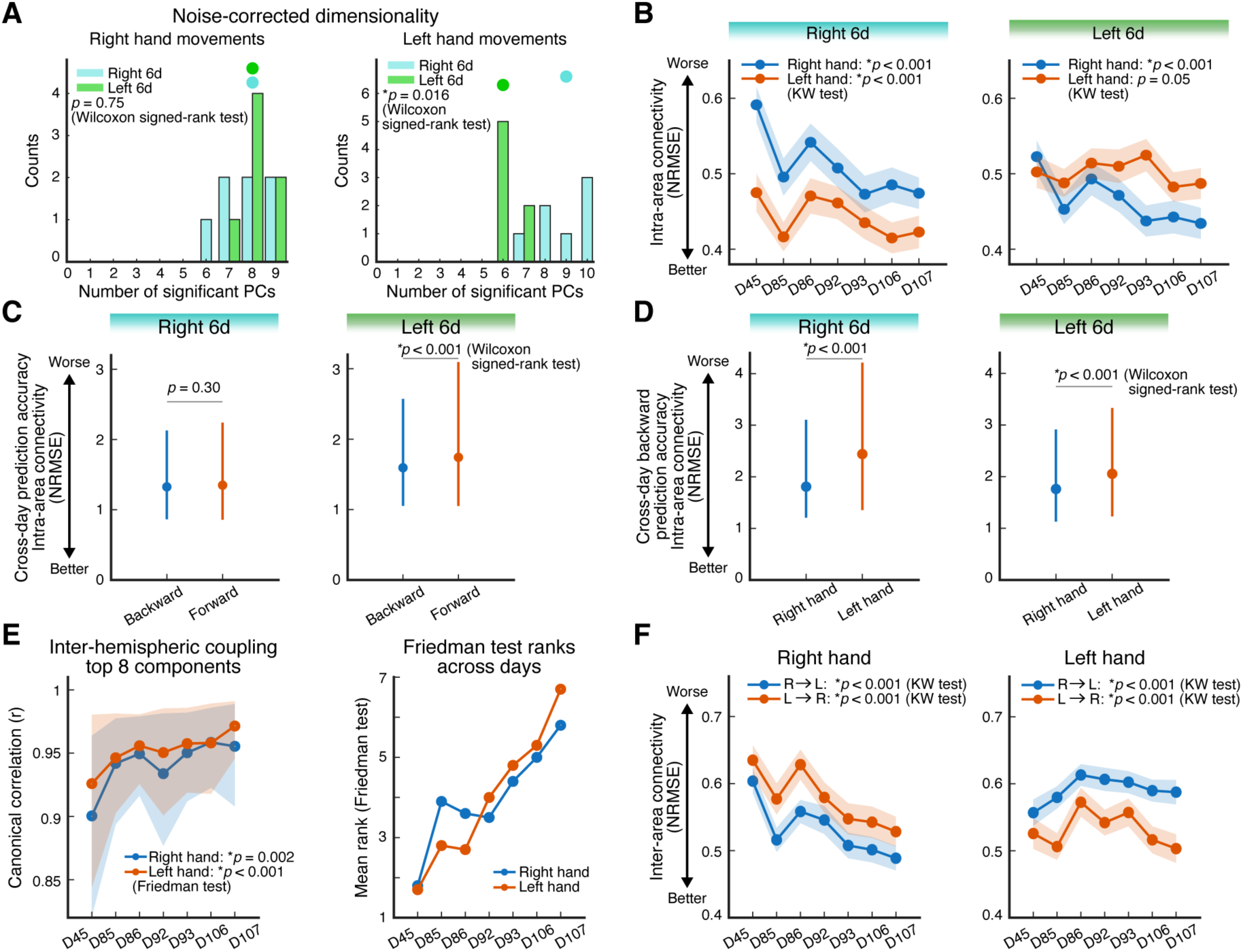
Session-wise intra- and inter-area connectivity dynamics in bilateral 6d during closed-loop finger typing. (A) Noise-corrected PCA dimensionality estimates for right 6d (cyan) and left 6d (green) across sessions, shown separately for right-hand and left-hand movements. Medians are indicated by dots above each distribution. For left-hand movements, dimensionality was significantly higher in right 6d (Wilcoxon signed-rank test, *p* = 0.016). (B) Intra-area functional connectivity evaluated using cross-validated GLM performance, quantified as normalized root mean squared error (NRMSE), for right and left 6d across sessions. An NRMSE value of 0 indicates perfect prediction (i.e., identical predicted and true signals), with larger values indicating poorer prediction performance. Both regions showed significant session-wise modulation for movements of the hand contralateral to each 6d (Kruskal–Wallis tests). (C) Cross-day GLM prediction performance within each area. “Forward” prediction trains on an earlier day and tests on a later day, whereas “Backward” prediction trains on a later day and tests on an earlier day. In right 6d, forward and backward predictions did not differ, whereas in left 6d, backward prediction showed significantly better performance (lower NRMSE) (Wilcoxon signed-rank test, *p<* 0.001). (D) Using backward prediction, right 6d and left 6d both showed significantly better performance for right-hand movements than for left-hand movements (Wilcoxon signed-rank tests, *p<* 0.001). Data are summarized as the median, with error bars indicating the 25th–75th percentiles (C, D). (E) Inter-area connectivity between right and left 6d estimated using CCA on the top eight PCA components (left panel). Canonical correlation showed significant session-wise modulation for both hands (Friedman tests: right hand *p* = 0.002; left hand *p <* 0.001). Session-wise Friedman mean ranks (right panel) indicate an overall increasing trend across sessions. (F) Inter-area connectivity across sessions. “R*→*L” indicates models trained on right 6d and tested on left 6d. All four conditions(hand *×* directional combinations) showed significant session-wise modulation (Kruskal–Wallis tests). Prediction tended to be better when decoding from right or left 6d during movement for the contralateral hand. In panels B, E, and F, data are shown as the mean, with shaded regions indicating the 95% confidence interval.

Importantly, for subsequent connectivity analyses, generalized linear models (GLMs) (Supplemental Figure S11) were not trained on raw channel activity. Instead, channels were first selected based on significant class-selective activity identified by FDR-corrected KW tests, and PCA was then applied to the selected channels. Based on the observed dimensionality estimates, a fixed number of components (top 10 PCs) was used as GLM input across all areas and sessions, with the aim of reducing potential confounds arising from differences in channel count between right and left 6d.

Connectivity was quantified by predicting each channel’s activity from all remaining channels within the same area using a leave-one-out PCA + GLM framework with 10-fold CV, and prediction error was quantified using normalized root mean squared error (NRMSE). An NRMSE value of 0 indicates perfect prediction, and larger values indicate poorer performance. In both 6d, intra-area connectivity within-session was overall better for contralateral than ipsilateral hand movements and varied significantly across days, with values tending toward lower NRMSE over time (Figure 6B).

To assess cross-session generalization, models were trained on one session and tested on either a later session (“forward”) or an earlier session (“backward”). In right 6d, prediction performance did not differ between forward and backward directions, whereas in left 6d, backward prediction performance exceeded forward prediction (Wilcoxon signed-rank test, *p<* 0.001; Figure 6C). This asymmetry indicates that the representational structure in left 6d became progressively more generalizable over days. Backward prediction also revealed better performance for right-hand movements than left-hand movements in both 6d (Figure 6D), suggesting greater representational stability for movements of the dominant (right) hand.

For inter-area connectivity, CCA was applied to the top eight PCs from each 6d (the smallest number of components capturing close to 90% of the shared variance, Supplemental Figure S12). Canonical correlations changed significantly across days and showed increasing ranks over time (Figure 6E). Finally, inter-area GLM prediction was evaluated by training on one area and predicting the other. Both prediction directions (R→L and L→R) showed significant cross-day changes (KW tests, *p <* 0.001). For both right- and left-hand movements, connectivity was consistently better when the prediction direction targeted the 6d during movement of the contralateral hand. Inter-are connectivity during right-hand movements showed lower NRMSE values in later sessions (Figure 6F).

### Compression of latent trajectories across sessions and distinct manifold structure across right and left 6d

To investigate changes in the low-dimensional space of relevant neural activity, dPCA was applied separately to each session with Hand, Finger, and Gesture marginalizations. Because not all finger tokens were extracted in every session following HMM labeling (Figure 2C), analyses were restricted to conditions shared across all sessions. This resulted in two Finger conditions—Thumb and Middle—which were the only fingers that appeared across all Hand and Gesture combinations in all sessions. For each marginalization, the top two dPCs were extracted, and trajectories were visualized in the dPC1–2 plane. Across days, latent trajectory magnitude decreased for all marginalizations (Hand, Finger, and Gesture) in both right and left 6d, with the reduction appearing visually most pronounced for Hand in right 6d (Figure 7A). Session-wise trajectory magnitudes decreased significantly for all marginalizations in both 6d (KW test, *p <* 0.001; Figure 7B), and magnitude was strongly negatively correlated with typing speed (CPM) but not with the RNN-derived CER (Figure 7C).

**Figure 7:**
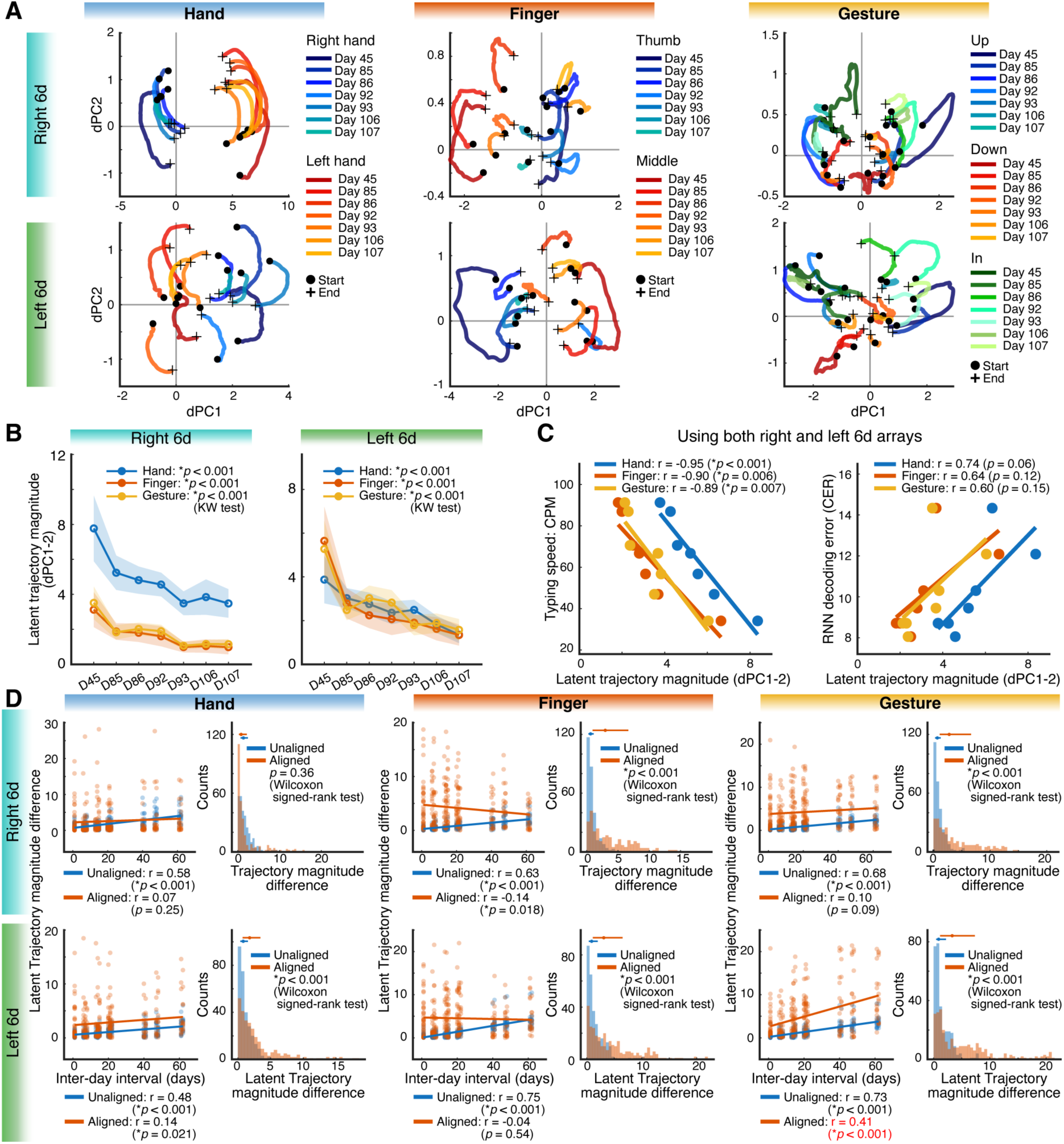
Session-wise changes in demixed neural trajectories during closed-loop finger typing. (A) dPCA was applied to HMM-segmented, z-scored FR from closed-loop typing sessions. For each marginalization (Hand, Finger, Gesture), neural trajectories were projected onto the first two demixed components (dPC1–2) after linearly resampling all trials to a common length of 100 samples. Trajectories begin at sample 1 (black dots) and end at sample 100 (crosses). Because not all tokens appeared in every session, Thumb and Middle were the only finger conditions present across all sessions. Across days, trajectory magnitude in the dPC1–2 plane progressively decreased. (B) Session-wise changes in trajectory magnitude for right and left 6d across the Hand, Finger, and Gesture marginalizations. All marginalizations in both 6d showed significant session-wise modulation (Kruskal–Wallis tests, *p<* 0.001) and exhibited decreasing trends. (C) Relationships between trajectory magnitude and behavioral indices. Typing speed (CPM) showed significant negative correlations with magnitude for all marginalizations, whereas RNN decoding error (CER) showed no significant correlations. (D) Cross-day differences in trajectory magnitude for right and left 6d. For unaligned dPC1–2 spaces (blue), the absolute magnitude difference increased with inter-day interval across all marginalizations (significant positive correlations). After CCA-based alignment (red), these positive correlations disappeared for all marginalizations in right 6d and for Finger in left 6d. Significant positive correlations remained for Hand and Gesture in left 6d. Histograms show the distributions of magnitude differences for unaligned (blue) versus aligned (red) spaces, with medians and the 25th and 75th percentile interquartile ranges shown above each panel. Magnitude differences were significantly larger in the aligned space than in the unaligned space for five of the six marginalizations (Wilcoxon signed-rank tests), except for the Hand marginalization in right 6d.

Because the dPC1–2 trajectory magnitude systemically decreased across sessions, larger inter-day intervals should produce larger differences in magnitude between two sessions. However, if learning-related changes are constrained by a manifold, CCA-based alignment should eliminate these interval-dependent differences ^13^. Consistent with this prediction, unaligned activity showed robust positive correlations between inter-day interval and trajectory-magnitude differences in all marginalizations (*p <* 0.001; Figure 7D, left panels). After CCA alignment, these positive correlations disappeared for all marginalizations in right 6d, consistent with learning-related changes that could be captured by CCA alignment. In contrast, in left 6d, alignment removed interval-related positive correlations only for Finger.

After alignment, observed negative correlations for right 6d Finger (*r* = −0.14) and positive correlations for left 6d Hand (*r* = 0.14) were nominally significant (*p* = 0.018 and 0.021, respectively); however, neither *p*-value would survive Bonferroni correction for six comparisons. In contrast, left 6d Gesture showed a robust positive correlation after alignment (*r* = 0.41, *p<* 0.001), indicating that the Gesture-related neural changes in left 6d were relatively resistant to CCA-based alignment.

## Discussion

In this study, we investigated how dorsal premotor neural population dynamics change during the acquisition of a new typing skill through repeated closed-loop iBCI use. Across sessions, representations in both right and left 6d became progressively more compact, as reflected by reduced Mahalanobis distances between finger-movement representations and contraction of low-dimensional neural trajectories in dPCA space. Importantly, trajectory magnitude was significantly negatively correlated with typing speed, but not with decoder performance, indicating that these changes reflected skill acquisition-related neural changes rather than increased separation of neural activity required for improvements in RNN decoder performance. Previous work has demonstrated that day-to-day changes in motor cortical population activity can be aligned across sessions using CCA, indicating that much of the neural variability may be captured within a neural manifold ^13^. Consistent with this observation, CCA-based alignment substantially reduced session-to-session differences in right 6d, suggesting that much of the observed neural changes may be representable within a low-dimensional space across sessions. In contrast, neural changes in left 6d were not fully captured by CCA alignment. Together, these results indicate that typing skill acquisition is associated with robust population-level compaction in premotor cortex, with common changes across bilateral 6d alongside selective changes expressed in left 6d.

Direct control of arm movements in the primate motor system is primarily mediated by contralateral descending projections ^29^. The present study demonstrated clear lateralization in FR modulation, with larger responses for contralateral than ipsilateral finger movements and prominent suppression in ipsilateral 6d during ipsilateral finger movements (Figure 1F), consistent with prior human reports ^30^. It should be noted that cortical excitability and interhemispheric inhibition can be altered following cervical spinal cord injury ^31^. Therefore, the activation and suppression patterns observed in T18 may not fully reflect premotor physiology in neurologically intact individuals.

More broadly, studies in non-human primates have shown that both primary motor cortex (M1) ^32^ and PMd ^33^ are engaged during contralateral as well as ipsilateral arm movements. Consistent with these findings, human neuroimaging studies have demonstrated that ipsilateral M1 and PMd exhibit task-related activity during both ipsilateral and contralateral finger movements ^30^. In addition, PMd has been implicated in movement planning and selection ^34–36^. While PMd involvement in motor output during limb movements has been widely reported, there has been little direct evidence in humans—particularly at the level of intracortical population activity—regarding how bilateral PMd representations differ during finger movements. Our population-level analyses revealed that right and left 6d did not encode finger-movement information in a mirror-symmetric manner. Instead, right 6d predominantly emphasized hand-related structure, whereas left 6d more strongly encoded Finger- and Gesture-related information (Figure 4C, D). In our clustering analyses, right 6d exhibited a clear separation between right- and left-hand movements, whereas such hand-based segregation was not evident in left 6d (Figure 1H, 3F, 5B). This pattern is consistent with prior reports of hemispheric asymmetry in PMd, in which left PMd has been implicated in movement selection for both hands, whereas right PMd is more strongly biased toward selection of contralateral (left-hand) movements ^37^. Together, these findings suggest that bilateral 6d supports finger movements through complementary representational roles rather than redundant encoding across hemispheres.

Previous positron emission tomography studies reported increased lateral premotor cortex activity during acquisition of novel motor sequences ^38^. Conversely, prior non-invasive studies have reported that practice is associated with reductions in task-related BOLD activity ^8,19,39,40^. Single-neuron population imaging studies have reported that learning is accompanied by a reduction in neural responsiveness and an increase in representational stability ^15^. At the level of intracortical population dynamics, we observed a complementary phenomenon: normalized Mahalanobis distances decreased across sessions, and the magnitude of low-dimensional neural trajectories in the dPC1–2 plane systematically declined over time. Together, these findings suggest that practice was associated with increasingly compact population-level neural activity, potentially enabling skilled neural typing to be performed with reduced neural load and greater efficiency.

At first glance, this result appears to contrast with prior motor learning studies reporting practice-related increases in representational dimensionality ^8^ and Mahalanobis distance ^23^, often accompanied by improved decoding accuracy. For example, Natraj et al. ^23^ showed that during learning of a novel BCI task, initially overlapping neural representations became progressively more separable, leading to increased Mahalanobis distances and improved decoder performance. In the present study, however, neural representations were already well separated at the outset (Figure 3C, 4G, H) and online decoder accuracy remained stable across sessions (Figure 2F). Under these conditions, practice was associated not with further representational separation but with a reduction in the magnitude of population-level neural trajectories. Together, these findings suggest that neural changes observed here reflect a learning process that differs from prior reports emphasizing increased representational separation with practice. Moreover, trajectory compression was strongly correlated with typing speed (CPM) but showed no relationship with decoder accuracy (Figure 7C), indicating that these changes primarily reflected modulation of the participant’s neural population dynamics rather than properties of the decoding algorithm.

CCA-based alignment demonstrated that learning-related changes in neural population dynamics during neural typing were not uniformly captured across right and left 6d. Prior work has shown that short-timescale BCI learning can be supported by changes occurring within a manifold ^9,10^. More recently, Oby et al. demonstrated in a non-human primate motor BCI task that long-term skill acquisition can involve the emergence of new neural population activity patterns, described as outside-manifold changes ^11^. While these studies directly assessed neural manifolds, Gallego et al. introduced an indirect approach using CCA-based alignment to assess neural non-stationarity in relation to within-manifold changes ^13^. In addition, day-to-day non-stationarity has been proposed to arise from drift along a neural manifold in prior population-level studies of motor behavior ^12,13,23,26,41,42^.

In the present study, CCA-alignment was used to compare neural population activity across sessions. Changes in right 6d were largely captured by alignment, consistent with learning-related changes occurring within a manifold. In contrast, gesture-related activity in left 6d was not well captured by alignment. Notably, left 6d also exhibited progressive increases in FR across sessions that were not observed in right 6d (Figure 5F), and forward-backward prediction analyses indicated cross-session generalization in left 6d (Figure 6C). Although the relationships among FR modulation, cross-session generalization, and alignment-resistant population changes were not directly tested, their co-occurrence supports that these phenomena may be linked during skill acquisition. Together, these results suggest that learning-related changes were more pronounced in left 6d, the dominant hemisphere, compared with right 6d. The inability to capture gesture-related learning in left 6d with CCA alignment raises the possibility that these changes may occur outside a manifold. However, because neural manifolds were not directly estimated in the present study, further work will be required to determine whether these changes truly represent outside-manifold learning.

Considering these results, left 6d may exhibit greater plasticity during skill acquisition than right 6d. In the BrainGate clinical trial, intracortical arrays have been implanted in the dominant hemisphere (typically, left), a strategy motivated by the goal of achieving robust and high-performance iBCI control ^43^. Prior work has suggested that, compared with right PMd, left PMd plays a more prominent role in movement selection and motor learning ^44–46^, and may serve as a hub within the motor learning network ^47^. Consistent with this framework, our results provide intracortical, population-level evidence supporting the rationale for implanting iBCI electrodes in the dominant hemisphere. At the same time, the robust typing performance achieved in this study likely depended on the complementary contributions of both hemispheres. Right 6d preferentially encoded hand-related information, whereas left 6d more strongly represented finger and gesture information, suggesting that bilateral recordings may provide richer neural information for high-performance typing iBCI.

Several limitations should be noted. This study is based on a single participant performing a closed-loop iBCI task. It is unknown whether similar population dynamics would be observed during natural typing practice in neurologically intact individuals or in non-BCI motor tasks. Although learning-related changes in right 6d were largely captured by CCA-based alignment, alignment revealed a significant negative relationship between neural trajectory magnitude and inter-session interval for Finger representations in right 6d, the interpretation of which remains unclear within the scope of the present analyses. More broadly, the associations between neural trajectory compression and typing speed are correlational rather than causal. In addition, interpretation of population structure may be influenced by asymmetries in electrode implantation: in the present study, dPCA results were highly consistent across the two arrays in right 6d and across the four arrays in left 6d, motivating aggregation within each hemisphere, but differences in array number and spatial coverage make it difficult to fully dissociate functional hemispheric asymmetries from effects related to electrode placement. Future studies across participants, implantation configurations, and task contexts will be needed to establish the generality and mechanisms underlying these effects.

In the low-dimensional population space, neural trajectories became progressively more compact as typing skill improved. Although this compression was observed bilaterally in 6d, across-day alignment revealed distinct learning-related dynamics across right and left 6d.

## RESOURCE AVAILABILITY

### Lead contact

Further information and requests for resources should be directed to and will be fulfilled by the lead contact, Daniel B. Rubin.

### Materials availability

This study did not generate new unique reagents.

### Data and code availability

The datasets generated and analyzed during this study are publicly available in the Dryad Digital Repository at https://doi.org/10.5061/dryad.cz8w9gjjk. The custom MATLAB code used for data analysis in this study will be made publicly available upon publication.

## ACKNOWLEDGMENTS

We thank T18, his family and care partners for the time and effort they contributed to this research. We thank Maryam Masood, Dave Rosler, and Beth Travers for their regulatory and team management efforts supporting this work. We thank Dr. Masayuki Hirata for providing support for the MATLAB license used in this work. This work was supported by: Office of Research and Development, Department of Veterans Affairs (A4820R), NIH NIDCD (U01DC017844, K23DC021297), NIH NINDS (U01NS123101), AHA (23SCEFIA1156586), the Japan Society for the Promotion of Science (JSPS Overseas Research Fellowships), and Japan-U.S. Brain Research Cooperative Program.

## AUTHOR CONTRIBUTIONS

H.H. developed the offline neural analysis pipeline, implemented all HMM-based labeling methods, generated all figures, and led the interpretation of results. H.H. wrote all MATLAB analysis code and contributed substantially to writing and revising the manuscript. J.J.J. H.L.A. and D.B.R. reviewed analysis and interpretation of experimental data. J.D.S. and J.J.J. installed hardware for neural recording with T18. J.J.J. collected T18 typing session data. J.D.S. managed integration and deployment of BCI system software and hardware. L.R.H. is the sponsor-investigator of the multisite BrainGate2 pilot clinical trial. Z.M.W. D.B.R. and L.R.H. planned T18’s array placement surgeries, and Z.M.W. performed T18’s array placement surgeries. L.R.H. was responsible for all clinical trial related activity at Massachusetts General Hospital (MGH). L.R.H. and D.B.R. supervised and guided all research activity with T18. The study was supervised and guided by D.B.R.. All authors reviewed and edited the manuscript.

## DECLARATION OF INTERESTS

CAUTION: Investigational Device. Limited by Federal Law to Investigational Use. The content is solely the responsibility of the authors and does not necessarily represent the official views of the National Institutes of Health, or the Department of Veterans Affairs, or the United States Government.

The MGH Translational Research Center has a clinical research support agreement (CRSA) with Ability Neurotech, Analog Devices, Axoft, Medtronic, Neuralink, Neurobionics, Paradromics, Precision Neuro, Reach Neuro, and Synchron, for which L.R.H. provides consultative input. Mass General Brigham (MGB) convenes the Implantable Brain-Computer Interface Collaborative Community (iBCI-CC); charitable gift agreements to MGB, including those received to date from Axoft, Blackrock Neurotech, Neuralink, Paradromics, Precision Neuro, and Synchron, support the iBCI-CC, for which L.R.H. provides effort.

## STAR*METHODS

### KEY RESOURCE TABLE

| REAGENT or RESOURCE | SOURCE | IDENTIFIER |
| --- | --- | --- |
| **Software and algorithms** |  |  |
| MATLAB | MathWorks | https://www.mathworks.com/ |
| FieldTrip toolbox | FieldTrip | https://www.fieldtriptoolbox.org/ |
| jPCA toolbox<br>(Churchland lab implementation) | Churchland laboratory | Churchland lab jPCA code |
| Tensor maximum entropy<br>(TME) | Elsayed and Cunningham, Nat Neurosci 2017 | https://github.com/gamaleldin/TME/ |
| demixed Principal Component Analysis<br>(dPCA) toolbox | Kobak et al., eLife 2016 | https://github.com/machenslab/dPCA/ |
| Piecewise Linear Time-Warping toolbox<br>(affinewarp) | Williams et al., Neuron 2020 | https://github.com/ahwillia/affinewarp/ |
| **Other** |  |  |
| NeuroPort Electrode | Blackrock Neurotech | https://blackrockneurotech.com/ |

### EXPERIMENTAL MODEL AND SUBJECT DETAILS

#### Human participant and ethics statement

This study was conducted under permission from the U.S. Food and Drug Administration and the Institutional Review Boards of Massachusetts General Hospital and Brown University. Research sessions were performed with one participant (T18), who was enrolled in the ongoing BrainGate2 clinical trial (ClinicalTrials.gov Identifier: NCT00912041). T18 was a right-handed man in his 40s with tetraplegia due to a cervical spinal cord injury (C4 AIS A) sustained approximately 18 months prior to enrollment. All research sessions were performed at the participant’s place of residence. This manuscript does not report clinical outcomes of the trial; instead, it presents scientific and engineering findings derived from neural and behavioral data collected within the context of the trial. T18 provided informed consent to participate in the study and consented to the publication of photographs and videos containing his likeness.

The BrainGate2 trial’s purpose is to collect preliminary safety information and demonstrate feasibility that an intracortical BCI can be used by people with tetraplegia for communication and control of external devices; the present manuscript results from analysis and decoding of neural activity recorded during the participants’ engagement in research that is enabled by the clinical trial but does not report clinical trial outcomes.

#### Electrode implantation

Before implantation, T18 underwent structural MRI, task-based functional MRI, and resting-state fMRI. These data were processed using the Human Connectome Project (HCP) ^1,2^ functional parcellation pipeline to guide identification of putative cortical targets for microelectrode array implantation. Based on this functional mapping, T18 received six 64-channel NeuroPort Electrode arrays (Blackrock Neurotech, Salt Lake City, UT; 1.5-mm electrode length); four implanted in the dorsal precentral gyrus (Brodmann 6d) of the left, motor-dominant hemisphere and two in the same region of the right hemisphere.

### METHOD DETAILS

#### Neural signal processing

Neural signal acquisition and preprocessing followed the standard BrainGate neural recording pipeline, which has been described in detail in previous publications ^3–5^. Accordingly, several aspects of the signal processing procedures described below resemble those reported in prior BrainGate studies.

Neural voltage time series signals were recorded and digitized (30 kHz, 16 bits per sample) using the Neuroplex-E system (Blackrock Neurotech) attached to three percutaneous connectors on each participant’s head, and transmitted via three mini-HDMI cables (one to each Neuroplex headstage), attached to two Gemini hubs (Blackrock Neurotech), prior to final processing via a Neural Signal Processor (NSP) (Blackrock Neurotech). Packets of neural data were streamed from the NSP to our processing pipeline. Signals were analog filtered (4th order Butterworth with corners at 250 Hz to 5 kHz) using the Scipy python library (scipy.signal.filtfilt).

Linear regression referencing ^6^ (LRR) filter coefficients and subsequent channel-specific thresholds were determined using the filtered digitized data recorded in an initial reference block at the beginning of each session. LRR coefficients are computed by solving *Y* = *WX* where *Y* is the signal from a given channel we require and *X* is the signal from all other channels. We solve for the LRR weight matrix through least squares calculation *W* : *W* = *inv*(*X^T^ X*)*X^T^ Y* where *inv* is matrix inversion ^7^. Channel-specific thresholds were calculated using filtered 30kHz data once these calculated references had been applied to identify spike events.

Thresholds were set at −3.5 times the standard deviation of the voltage signal for each channel. Neural features were computed at 100 Hz (10 ms bins). Specifically, the number of non-causal threshold crossings ^8^ (ncTX) was obtained by counting threshold crossings within each 10 ms bin of the filtered neural signal, and spike-band power (PW) was computed as the sum of squared voltages within the same 10 ms bin. During closed-loop decoding blocks, feature normalization was employed to account for neural non-stationarities (drifts in mean firing rate) which could arise over the course of a block. Within each channel, ncTX rates and PW were z-scored (mean subtracted and divided by standard deviation per channel). Feature extraction (ncTX and PW), binning, decoding and task phase control were performed through the Python based, modular BRAND ^9^ framework, where each process is instantiated as a self-contained node-based Python program. Messaging between these nodes is performed using a Redis database.

#### Finger Movement task

In this instructed-delay paradigm, each trial for T18 began with a preparatory period that varied between 1.5 and 2.5 seconds. This was followed by a 2.5-second movement period (including a required hold) and a 2.5-second rest period (Figure 1B). Throughout the preparatory phase, the participant was instructed to remain in the rest posture until the Go, indicated by the on-screen rectangle turning green accompanied by a beep tone. At the Go cue, T18 was instructed to attempt and hold one of the three active gestures associated with a specified finger (Figure 1C) until the “Return” cue appeared, accompanied by a click sound. This task included 30 finger movement conditions, defined by a factorial combination of Hand (right or left), Fingers (five digits), and Gesture (Up, Down, In). In addition to these movement trials, a “Do Nothing” condition (denoted as “None” in the manuscript) instructed the participant to remain at rest posture and perform no intended movement throughout the entire Go period.

#### Neural separability during the Finger Movement task

To assess the degree of neural separability across finger movements, we analyzed neural activity recorded during the open-loop Finger Movement task. The following analyses quantify how distinctly different finger-movement conditions were represented in population neural activity.

#### Cluster-based spatiotemporal permutation test of the Finger Movement task

Neural activity acquired during the Finger Movement task was quantified as Gaussian-smoothed threshold crossing rates derived from ncTX. ncTX were converted to continuous rate estimates by convolution with a Gaussian kernel (*σ* = 30 ms), corresponding to three 10-ms time bins. Throughout the manuscript, these smoothed threshold crossing rates are referred to as firing rates (FR).

To identify task-related neural responses, FR during the active phase following the Go cue were compared with activity during the preparation phase. For each trial, baseline activity was computed by averaging FR across the preparation phase and replicated across the active time window to match the temporal structure of the active data.

Statistical comparisons were performed independently for each task. For each channel and time point, paired-sample t-statistics were computed across trials (*n* = 10), comparing activity during the active phase with the corresponding baseline. To correct for multiple comparisons across channels and time, statistical significance was assessed using a cluster-based spatiotemporal permutation test ^10^, as implemented in FieldTrip ^11^.

Clusters were defined based on spatial adjacency between neighboring channels and temporal contiguity, using a cluster-forming threshold of *p <* 0.05. Cluster-level statistics were computed as the sum of t-values within each cluster (maximum cluster sum). Statistical significance was determined by comparing observed cluster statistics against a nonparametric null distribution generated via 1,000 Monte Carlo permutations. All statistical tests were two-sided with a final significance threshold of *α* = 0.05. Statistical analyses were performed using the `ft_timelockstatistics` function in FieldTrip.

Spatial adjacency was defined based on the physical 8 × 8 NeuroPort electrode geometry for each subarray, accounting for array-specific orientation. Neighboring channels were identified using distance-based criteria implemented in FieldTrip.

#### Quantification and visualization of significant time points

To summarize the spatiotemporal cluster results at the level of individual channels, we quantified the temporal extent of significant task-related activity for each channel. For each channel, we counted the number of time points that were identified as significant within any spatiotemporal cluster within a predefined post–Go presentation window (1–100 data points, corresponding to 0–1 s after the Go presentation).

This count was used as a measure of the duration of significant modulation and was visualized as the size of the channel marker in spatial maps. Channels exhibiting exclusively positive t-values across significant time points (increased FR relative to baseline) were shown in red, whereas channels exhibiting exclusively negative t-values (decreased FR relative to baseline) were shown in blue. Channels that exhibited both positive and negative significant t-values within the same time window were shown in green; such channels were rare (Figure 1D, Supplemental Figure S1).

#### Validation of spatiotemporal cluster analysis using PSTHs

To assess the validity of the spatiotemporal cluster analysis, FR were averaged across trials (n = 10) to generate peristimulus time histograms (PSTHs). In Figure 1E, we show two representative channels in right 6d during the right middle finger upwards condition: Ch 374, which exhibited a significant positive t-value, and Ch 360, which exhibited a significant negative t-value.

Consistent with the cluster-based statistical results, Ch 374 showed a clear increase in FR around the Go presentation, whereas Ch 360 showed a suppression of FR over a similar time period. In addition, for Ch 360, PSTHs are shown for both right- and left-hand movements, demonstrating that the direction of firing rate modulation differed between right- and left-hand movements.

Together, these examples confirm that positive t-values identified by the spatiotemporal cluster analysis correspond to increases in FR (activation), whereas negative t-values correspond to decreases in FR (suppression).

For completeness, PSTHs computed from ncTX and PW are shown in Supplementary Figure S2. Compared to these representations, PSTHs based on FR exhibit more clearly defined temporal profiles, facilitating visual interpretation of condition-dependent modulation.

#### Temporal dynamics of normalized mean t-values

Temporal dynamics of activation and suppression were quantified by grouping spatiotemporal cluster statistics by movement sets. Finger movement conditions were grouped into right-hand and left-hand movements (15 conditions each). For right and left 6d, spatiotemporal cluster results were pooled separately for right-hand and left-hand finger movement groups and analyzed separately for activation (*t >* 0) and suppression (*t <* 0).

For each time point, we computed the proportion of significant channels by dividing the number of significant channels by the total number of channels in the corresponding array (128 channels for right 6d and 256 channels for left 6d). We then summarized the distribution of significant t-values across channels by excluding non-significant channels and estimating mean t-values separately for activation and suppression. To account for time-varying recruitment of significant channels, we additionally computed a normalized mean t-value by weighting the mean t-value at each time point by the corresponding proportion of significant channels, and estimated the mean and 95% confidence interval using `normfit` (MATLAB).

The resulting activation and suppression time courses were smoothed using Gaussian kernel filtering (*σ* = 0.02 s). Normalized mean t-values were computed at each time point aligned to the Go presentation (*t* = 0) and plotted over time (Figure 1F).

#### Spatial similarity of task-evoked activity patterns using Hamming distance

Spatial changes in task-evoked activity patterns were quantified by comparing channel-wise patterns of significant modulation across finger movement conditions. For each movement condition and for each area (right 6d and left 6d), within a 1-s time window, each channel was assigned a categorical label based on the direction of significant modulation identified by the spatiotemporal cluster analysis: 0, no significant modulation; 1, significant activation only (*t>* 0); 2, significant suppression only (*t<* 0); and 3, both significant activation and suppression observed within the same time window.

For each pair of movement conditions, we computed a masked Hamming distance between the corresponding channel-label vectors. Channels were included in the comparison if they were nonzero in either condition (i.e., at least one condition exhibited significant modulation at that channel). The Hamming distance was defined as the fraction of included channels whose labels differed between the two conditions, yielding values between 0 (identical spatial labeling patterns) and 1 (maximally different patterns). Distances were computed separately for right 6d (128 channels) and left 6d (256 channels).

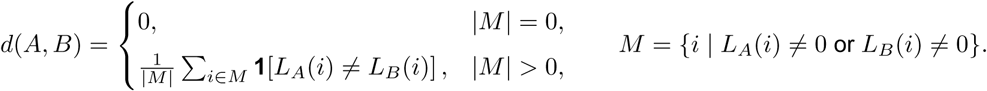

where *A* and *B* denote two movement conditions, and *L_A_*(*i*) and *L_B_*(*i*) are the categorical labels assigned to channel *i* for conditions *A* and *B* (0: none, 1: activation only, 2: suppression only, 3: both). *M* = *{i | L_A_*(*i*) ≠ 0 or *L_B_*(*i*) ≠ 0*}* is the set of channels showing significant modulation in at least one condition, and **1**[*·*] is the indicator function. If *|M|* = 0, we defined *d*(*A, B*) = 0.

Distances were computed across sliding time windows throughout the trial; however, Figure 1G focuses on the 0.2–1.2 s window, which exhibited the clearest task-dependent spatial differentiation.

#### Hierarchical clustering and dendrogram visualization

Dendrograms were constructed to visualize the similarity structure across finger movement conditions using task-by-task distance matrices derived from masked Hamming distances (Figure 1H) or normalized Mahalanobis distances (Figure 3F, Figure 5B). The symmetric distance matrix was converted to a vector containing all unique pairwise distances (MATLAB `squareform`). Hierarchical clustering was then performed using agglomerative clustering with Ward linkage (MATLAB `linkage`), and dendrograms were plotted with all leaves shown (MATLAB `dendrogram`). For visualization, leaf labels were color-coded according to predefined movement groupings (e.g., right/left hand or up/down/in).

We compared dendrograms constructed using average and Ward linkage and observed no substantive differences in the overall clustering patterns (Supplementary Figure S3). Because the dendrograms were used for visualization, we report results using Ward linkage, as it provided visually clearer dendrograms.

#### Selection of class-selective channels

To identify channels exhibiting class-selective neural activity, we evaluated neural responses across all 31 conditions (30 finger movements plus a “None” condition) on a per-channel basis. For each channel, task-related differences in neural activity were assessed using a nonparametric Kruskal–Wallis (KW) test across conditions.

Analyses were performed separately for three neural features: z-scored ncTX (z-ncTX), z-scored PW (z-PW), and FR. For z-ncTX, the mean ncTX value was subtracted separately within each recording block, and the mean-centered ncTX values were then pooled across blocks to estimate a global standard deviation. The centered ncTX signals were divided by this global standard deviation to obtain z-ncTX. z-PW signals were computed using the same procedure. FR was computed from ncTX using a Gaussian kernel (*σ* = 30 ms).

For each feature, statistical testing was conducted at each time point relative to the Go. Channels showing significant task-dependent modulation were identified and referred to as class-selective channels. The total number of significant channels was quantified as a function of time around Go (total channels = 384; 6 arrays × 64 channels).

To account for multiple comparisons across channels, false discovery rate (FDR) correction was applied with a significance threshold of *α* = 0.05. The temporal evolution of the number of significant channels for each neural feature is shown in Figure 3A.

#### SVM decoding of Finger Movement conditions

To quantify time-resolved decoding performance across the 31 conditions (30 finger movements plus “None”), we trained a multi-class linear support vector machine (SVM) classifier using z-ncTX and z-PW features. Decoding was performed in sliding time windows aligned to the Go cue. Neural data were segmented from −1.5s to after 2.4 s around the Go. A 1.0-s analysis window (100 bins) was advanced in 0.1-s steps (10 bins), and each window was assigned a center time relative to Go.

For each time window, input features were restricted to class-selective channels identified at the same window using the KW+FDR procedure described above. Channel selection was performed separately for z-ncTX and z-PW; for each feature, only channels passing FDR (*α* = 0.05) were retained. These two features were used for decoding because they yielded a larger number of class-selective channels than FR (Figure 3A). The final input vector for each trial consisted of the concatenation of the selected z-ncTX and z-PW channels.

Decoding was evaluated using stratified 10-fold cross-validation. Within each fold, feature standardization was performed using training data only: for each channel/feature, the mean and standard deviation were computed from the training trials (pooling all time points within the current window), and both training and test data were standardized using these training-derived parameters. Standardized signals were then averaged across time within the window, yielding one feature vector per trial. A multi-class linear SVM was trained using MATLAB’s `fitcecoc` function. Predictions were generated for held-out trials in each cross-validation fold. Decoding accuracy was computed as the proportion of correctly classified trials aggregated across folds, and plotted as a function of time relative to Go (Figure 3B).

To assess anatomical contributions, the same decoding pipeline was applied using channels from (i) all arrays, (ii) right 6d (channels 257–384), and (iii) left 6d arrays (channels 1–256), with class-selective channel selection performed within each subset.

#### MLP decoding

As in the SVM decoding, a combination of z-ncTX and z-PW features was used to train the multi-layer perceptron (MLP) model, and only class-selective channels for each feature were included as inputs.

Decoding performance was evaluated using 5-fold stratified cross-validation. In each fold, the held-out 20% of the data were split evenly into validation and test sets (10%/10%), and this validation/test assignment was swapped once per fold (i.e., two evaluations per fold). Feature normalization parameters (mean and standard deviation) were computed using the training set only and applied to the validation and test sets. After z-scoring, feature vectors were optionally L2-normalized on a per-trial basis.

To improve robustness, we augmented the training data by fitting a class-conditional multivariate Gaussian to the training samples of each class (class-specific mean and covariance). For each class, we generated *M* synthetic samples per original training sample (*M* = 5) by sampling from the fitted Gaussian (with a small diagonal jitter added to the covariance for numerical stability). The synthetic samples were concatenated with the original training set and shuffled prior to training.

MLP classifiers were implemented in MATLAB using the `trainNetwork` function with a `featureInputLayer`. The network consisted of a single hidden layer with 165 units, followed by a softmax output layer. Leaky ReLU activations (*slope* = 0.01) and dropout (*rate* = 0.4) were applied to the hidden layer. Training used the Adam optimizer with a mini-batch size of 32, early stopping based on validation performance (patience = 5 validation checks), and a piecewise learning-rate schedule (drop factor of 0.5 every 10 epochs). This architecture was chosen based on prior work ^12^. Using a 1-s time window centered at 0.7 s after the Go, the MLP achieved a decoding accuracy of 94.2% (Figure 3C).

#### Latent-space analysis with an autoencoder

To visualize the low-dimensional structure of neural population activity, we trained a supervised autoencoder in which the 2D latent space was jointly constrained by reconstruction and task-classification objectives. Input features were identical to the windowed feature vectors used for MLP decoding. This approach follows established methods described in prior work ^12^.

The encoder consisted of two fully connected (FC) layers with ELU activation and dropout (*rate* = 0.3):

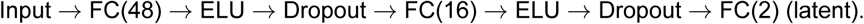

From the 2D latent layer, the network branched into two heads:

1. Decoder: FC(16) → ELU → FC(48) → ELU → FC(InputSize), which reconstructed the input features.
2. Classifier head: FC(#classes) → softmax, which predicted the movement class from the latent representation.

Training minimized a weighted sum of reconstruction error (mean squared error) and classification loss (cross-entropy), with a mixing weight of *α* = 0.5, optimized using the Adam optimizer. After training, latent coordinates were extracted from the 2D latent layer for visualization (Figure 3D).

For visualization, class-specific ellipses were overlaid in the latent space. For each class, an ellipse was derived from the covariance of its latent points by eigendecomposition of the 2 × 2 covariance matrix, scaling a unit circle (*scalefactor* = 2), and translating it by the class mean.

#### Mahalanobis distance

To quantify condition separability directly in neural feature space, we computed pairwise Mahalanobis distances among the 31 Finger Movement conditions (30 finger movements plus a None condition) using z-ncTX and z-PW features. For each array group (right 6d, left 6d, and all arrays), all available channels were included, and z-ncTX and z-PW were concatenated to form the input feature vector. Neural activity was analyzed in 1-s sliding windows (100 bins at 10-ms resolution) spanning −0.5 to 1.5 s relative to the Go, with a step size of 0.1 s. Within each time window, neural activity was averaged across time, yielding a Task × Trial × feature representation. The dimensionality of the input features depended on the number of channels (e.g., right 6d: 64 channels per array × 2 arrays × 2 features = 256; left 6d: 64 channels per array × 4 arrays × 2 features = 512).

For each condition, we computed the mean feature vector across trials and the trial-to-trial covariance matrix. The Mahalanobis distance between conditions *i* and *j* was defined as the distance between their mean vectors using a pooled covariance matrix, computed as the average of the two condition-specific covariance matrices. To ensure numerical stability, Tikhonov regularization was applied by adding a small diagonal term (*λI*, with *λ* = 0.001) to the pooled covariance matrix. This procedure yielded a 31 × 31 distance matrix for each time window and array group. Pairwise Mahalanobis distance between task-averaged population vectors was computed as:

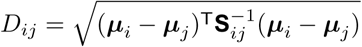

where ***µ****_i_* and ***µ****_j_* denote the mean feature vectors for tasks *i* and *j*, respectively.

The pooled covariance matrix **S***_ij_* was defined as:

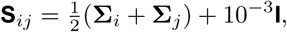

where **Σ***_i_*and **Σ***_j_*are the trial-wise covariance matrices for tasks *i* and *j*, and **I** is the identity matrix.

Because Mahalanobis distance scales with the dimensionality of the input space, distances were normalized by the square root of the number of input features. All reported values and visualizations therefore use this dimension-normalized Mahalanobis distance. For each time window, overall separability was quantified by pooling Mahalanobis distances across all unique condition pairs (31 choose 2) and computing their mean. Ninety-five percent confidence intervals were estimated using a normal approximation applied to the pooled distribution of pairwise distances (Figure 3G).

To visualize the structure of condition relationships, we performed hierarchical clustering and constructed dendrograms (see “Hierarchical clustering and dendrogram visualization”). The distance matrices (Figure 3E) and dendrograms (Figure 3F) correspond to the 1-s window centered at 0.7 s after the Go, which coincided with the peak SVM decoding accuracy.

To relate geometric separability to decoding performance, we paired the time-resolved mean normalized Mahalanobis distance with the corresponding time-resolved SVM decoding accuracy across matched windows and quantified their association using correlation analysis and linear regression (Figure 3H). For left 6d, we additionally evaluated dorsal and ventral subgroups (two arrays each), as these subgroups exhibited markedly different decoding accuracies.

#### jPCA of the Finger Movement task

To quantify low-dimensional rotational dynamics shared across finger-movement conditions, we applied jPCA using the publicly available implementation from the Churchland laboratory ^13^. Neural activity during the Finger Movement task was analyzed using all recorded channels and all movement conditions (30 conditions; the “None” condition was excluded from jPCA).

Analyses were performed on FR derived from ncTX and z-scored relative to the “None” condition, as described below. ncTX were converted to FR by convolution with a Gaussian kernel (*σ* = 30 ms) ^14^. ncTX recorded during the None condition were processed identically, and the mean and standard deviation of the resulting FR were computed across all None trials. FR for each finger-movement condition were then z-scored using this None-derived mean and standard deviation, yielding z-FR.

For preprocessing prior to jPCA, PSTHs were computed from z-FR for each condition. For each channel, PSTHs were soft-normalized by dividing by the response range computed across all conditions and time points plus a constant (*α* = 10), and then mean-subtracted across conditions at each time point. Dimensionality reduction was performed using PCA, retaining the top six principal components (*pcaDim* = 6). jPCA was then applied to the PCA-reduced trajectories over specified analysis intervals.

To identify time periods exhibiting strong rotational structure, we performed a grid search over (i) the center time of the analysis window and (ii) the window length. Window lengths ranging from 200 to 600 ms were tested in 100-ms increments. For each window length, the window start time was stepped from −500 ms to +1000 ms in 10-ms increments, defining target ranges [tStart, tEnd] in milliseconds. For each combination of window length and time position, jPCA was fit using identical preprocessing and PCA settings. Results for analysis windows spanning −0.2 to 0.6 s relative to the Go are shown in Supplemental Figure S4A.

Rotational dynamics were quantified using the goodness-of-fit of the best two-dimensional skew-symmetric dynamical system in the jPCA plane, reported as the *M*_skew_ *R*^2^ (Supplemental Figure S4A). This metric captures how well a purely rotational model explains the temporal derivatives of population activity projected into the jPCA1–2 plane.

To assess the statistical significance of the observed rotational dynamics, surrogate datasets were generated using a tensor maximum entropy (TME) approach ^14,15^. Based on the grid search, strong rotational structure was observed for a 200-ms window centered at 0.25 s relative to the Go. Statistical testing was therefore performed for this analysis window. For visualization, population trajectories for each finger-movement condition were projected onto the first two jPCA axes and plotted over time, with trajectories traced across the analysis window in the jPCA1–2 plane (Supplemental FigureS4B).

We constructed a tensor of mean-subtracted, soft-normalized PSTHs arranged as time × channels × conditions (finger movements). A TME model was fit to these data, constrained to match the marginal covariances across time, channels, and conditions, as well as the mean tensor. From the fitted model, 1,000 surrogate tensors were sampled. Each surrogate dataset was processed through the same PCA (top six PCs) and jPCA pipeline, and the two-dimensional *M*_skew_ *R*^2^ was computed. Statistical significance was assessed using an empirical one-sided p-value:

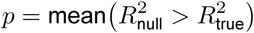

where 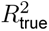 is the observed value and 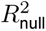 denotes the surrogate distribution (Supplemental Figure S4C).

#### dPCA of the Finger Movement task

Neural activity recorded during the open-loop Finger Movement task was analyzed using demixed principal component analysis (dPCA). ncTX were converted to FR by convolution with a Gaussian kernel (*σ* = 50 ms) ^16^. ncTX recorded during the None condition were processed in the same manner, and the mean and standard deviation of the resulting FR were computed across all None trials. FR for each finger movement condition were then z-scored using this None-derived mean and standard deviation, yielding z-FR.

To perform dPCA, marginalizations corresponding to task structure were defined. The 30 finger movement conditions consisted of a factorial combination of Hand (right or left), Finger (thumb, index, middle, ring, pinky), and Gesture (Up, Down, In). Accordingly, Hand, Finger, and Gesture were specified as task-related marginalizations. Activity that was shared across conditions and varied primarily with time was designated as the Common marginalization, while variance not accounted for by these terms was assigned to Interactions.

dPCA was performed using publicly available implementations. Regularization strength was selected using `dpca_optimizeLambda` (*numRep* = 5 and `simultaneous` = true), and noise covariance was estimated using `dpca_getNoiseCovariance` (`simultaneous` = true). dPCA was then performed using the `dpca` function with the optimized regularization parameter and the estimated noise covariance, extracting 20 components under the predefined marginalization structure. Variance explained by each component and marginalization was quantified using `dpca_explainedVariance`.

Cumulative explained variance for PCA and dPCA was computed and visualized (Figure 4A, B, top). In addition, the contribution of each marginalization to individual dPCA components was quantified (Figure 4A, B, bottom). For each component, the marginalization explaining the largest fraction of its variance was identified, and that marginalization was assigned as the representative label of the component. For example, because the first dPCA component in the right 6d array was primarily explained by Hand, it was labeled as the Hand dPC1. Using this procedure, all 20 components were labeled, resulting in each marginalization being represented by multiple components. The total variance explained by each marginalization across components was summarized using pie charts (Figure 4C, D).

In addition, pie charts summarizing the total variance explained by each marginalization were generated separately for each individual electrode array and are shown in the Supplemental Figure S7. These array-level analyses revealed highly consistent marginalization profiles across the two arrays implanted in right 6d and across the four arrays implanted in left 6d. Based on this consistency, and to facilitate interpretability of population-level results, dPCA analyses were performed by pooling arrays within each hemisphere. Specifically, data from the two right 6d arrays were combined and treated as representative of right 6d, and data from the four left 6d arrays were combined and treated as representative of left 6d. This operational definition enabled direct comparison of population-level dynamics between right and left 6d, while acknowledging that effects of array number and spatial coverage cannot be fully dissociated from functional hemispheric differences. This same pooling strategy was applied consistently across all subsequent analyses when referring to right and left 6d.

To visualize temporal dynamics, regularized projection matrices (*Wr*) obtained from the regularized dPCA were applied to trial-averaged PSTHs for each task, yielding time-resolved dPC trajectories. For each marginalization, the top-ranked component (i.e., the component explaining the largest fraction of variance for that marginalization) was selected, and its time course was plotted for all task conditions (Figure 4E, F).

Next, for each marginalization, up to three dPCA components labeled with that marginalization were selected, and PSTHs were projected onto the resulting dPCA space. Depending on the number of available components, trajectories were visualized in one-, two-, or three-dimensional dPCA representations (Figure 4G, H). For Gesture-related activity, the right 6d array contained two gesture-related components and was therefore visualized in a two-dimensional plane, whereas the left 6d array contained three such components and was visualized in three-dimensional space.

All analyses were performed separately for the right and left 6d arrays. Neural activity from −1.5 s to 2.4 s relative to Go presentation was analyzed. In dPC trajectory plots, the starting time point (−1.5 s) is indicated by filled circles, and the ending time point (2.4 s) is indicated by plus symbols (Figure 4G, H).

#### Online decoder outputs used for closed typing analysis

In the closed-loop typing task, T18 was instructed to copy type sentences presented on the screen. To implement the closed typing task, the 30 finger movements used in the previous paradigm were mapped onto individual keys arranged in a QWERTY-style keyboard layout (Figure 2A). T18 was instructed to perform the copy-typing task as if touch typing, using attempted finger movements to select each key. Although he was initially unfamiliar with touch-typing, his performance improved noticeably over successive sessions.

We analyzed neural and behavioral data collected during this closed-copy-typing session. The online decoder used in this task consisted of a 5-layer gated recurrent unit (GRU) recurrent neural network (RNN) trained with Connectionist Temporal Classification (CTC) loss to output character probabilities. A 5-gram language model implemented using a weighted finite-state transducer (WFST) ordinarily converts these probabilities into predicted words. All 384 channels were used, and two neural features—ncTX and spike-band power (PW)—were extracted from each channel, yielding a total of 768 input features to the RNN (binned in 20-ms windows). Character probabilities were inferred every 120 ms using a 300-ms window of preceding neural activity.

These decoding methods, including the full RNN architecture, training procedures, and language model implementation, were developed within the BrainGate2 framework and have been described extensively in prior publications ^4,5,17^. We refer readers to those reports for comprehensive methodological details and do not repeat them here.

Importantly, for all analyses in this manuscript, we treated the RNN-decoded character probabilities as the participant’s intended typing behavior and did not use the final outputs of the 5-gram language model. Accordingly, we evaluated decoding performance solely using the raw RNN predictions (top-1 outputs) and computed character error rate (CER) based on these predictions.

#### Preparation of neural templates for HMM-based forced-alignment labeling

During closed-typing sessions, neural activity was streamed online into an RNN, which output a probability distribution over 31 token classes: 30 character tokens corresponding to distinct finger movements and a blank state. Although the RNN provides a time series of estimated probabilities for each token based on past neural activity, its outputs reflect temporally integrated information rather than the instantaneous onset of the participant’s intended movement. As a result, the precise timing at which the participant (T18) attempted the corresponding keystroke cannot be directly inferred from the RNN output alone.

To resolve this uncertainty, we performed offline labeling of the recorded neural data to estimate the timing of attempted keystrokes. We adapted a forced-alignment labeling approach based on hidden Markov models (HMMs), following prior work in neural decoding of handwriting ^3^, and modified it to accommodate distinct finger-movement-based character tokens. This procedure enabled more accurate estimation of the start and end times of each inferred token.

#### Template construction for forced-alignment labeling using an HMM

Neural templates for forced-alignment labeling were constructed using data from an open-loop finger movement task (see “Finger Movement task”). ncTX signals were z-scored using the same procedure described in the Selection of class-selective channels. z-ncTX signals were converted to FR by convolution with a Gaussian kernel (*σ* = 30 ms) ^3^, yielding z-scored FR (z-FR). For each finger movement condition, z-FR data were segmented around the Prepare and Go presentations, producing trial × time × channel matrices.

To account for variability in movement timing across trials, trial-wise neural activity for each finger movement condition was temporally aligned using a piecewise time-warping procedure ^18^. The piecewise linear time-warping toolbox (https://github.com/ahwillia/affinewarp), implemented in Python, was used, and we applied the `PiecewiseWarping` model provided by this toolbox. For each condition, the `PiecewiseWarping` model was fit to the z-FR data (trials × time × channels) using one knot (*n* − *knots* = 1), a warp regularization scale of 0.001 (L1), and a smoothness regularization scale of 1.0 (L2). Model fitting was performed with 50 outer iterations and 200 warp-optimization iterations per outer iteration (Supplemental Figure S5A).

After time-warping, the aligned activity was averaged across trials for each condition to obtain a condition-specific mean response (PSTH; time × channels). To define the template duration for each finger-movement token, we inspected the root-mean-square (RMS) amplitude of the PSTH across channels and manually selected an end time for each condition. Specifically, the end time was defined as the point at which the RMS activity returned to within ±0.5 SD of the pre–Go baseline and remained near baseline for approximately 200–300 ms. End points were determined by visual inspection based on this criterion (Supplemental Figure S5B).

For each condition, we then excluded the first 100 ms after the Go presentation to remove early post-cue activity, and extracted the PSTH segment from 0.1 s to the condition-specific end time. The resulting PSTH segment (10-ms resolution) was down-sampled to 50-ms resolution by averaging consecutive non-overlapping 5-bin windows. The final 50-ms templates for all 31 tokens (30 finger-movement character tokens plus the blank/none token) were saved in MATLAB and used to construct HMMs for forced-alignment labeling.

#### HMM-based forced-alignment labeling without Blank for template refinement

To address session-to-session and within-session nonstationarity in neural activity, neural templates constructed from the open-loop Finger Movement task were further refined before final forced-alignment labeling. Because neural representations during closed typing gradually drifted over time, templates derived from the initial open-loop data alone became progressively less well matched to the closed-typing data. To mitigate this mismatch, we first performed HMM-based forced-alignment labeling (HMM labeling) without a Blank state (i.e., excluding the “none” template). This refinement stage used only the 30 finger-movement templates derived from the Finger Movement task and served to adapt the templates to session-specific neural activity before introducing a Blank state in the final 31-class HMM.

##### Preprocessing of typing-session neural activity

For each closed copy-typing session, we extracted trial-wise ncTX during the post–Go period for each prompted sentence. In this task, after T18 finished copying a prompted sentence, a 3-second instructed rest period was inserted before the next sentence began.

Within each session, ncTX was converted to FR by convolving each channel’s 10-ms binned spike counts with a Gaussian kernel (*σ* = 40 ms) ^3^ using reflect padding to reduce edge artifacts, and scaling to Hz. FR was then z-scored using statistics computed from the same session’s rest periods. Specifically, we concatenated neural activity during all instructed rest epochs, computed the session-level mean and standard deviation for each channel, and used these values to z-score the FR during typing epochs.

##### Building a left-to-right HMM from movement templates and ground-truth text

For each sentence, we tokenized the ground-truth transcription and mapped each token to an index in the keyboard label set; tokens absent from the template set were excluded. Using the resulting token sequence, we constructed a left-to-right HMM in which each token was represented by a chain of states whose emission means were defined by the corresponding template time bins (i.e., each template time bin constituted one HMM state).

The HMM comprised (i) an initial state distribution fixed to the first state of the first token; (ii) a sparse transition matrix that allowed within-token forward progression with limited skipping (stay/advance/skip-by-2); and (iii) a state-wise emission mean matrix *µ* assembled by concatenating the 50-ms template trajectories corresponding to the token sequence. For intermediate states within a token, self, next-state, and skip-by-2 transitions were permitted with probabilities on the order of 0.2, 0.6, and 0.2, respectively.

Emission likelihoods were computed under an isotropic Gaussian model with covariance *σ*^2^*I* (*σ*^2^ = 1, consistent with z-scored observations), yielding a time-by-state log-likelihood matrix.

##### Temporal constraints and Viterbi decoding

To stabilize forced alignment, we imposed two constraints on the emission log-likelihood matrix prior to decoding, following the forced-alignment strategy used in Willett et al. (2021) ^3^. First, for each token, we restricted allowable states to occur within a broad time window centered at the token’s expected relative position in the sentence (±30% of the sentence duration). Second, we enforced a terminal constraint such that the final frame could only occupy the final state of the final token. We then decoded the most likely state sequence using a Viterbi algorithm in log space (Supplemental Figure S5C).

The decoded state path was converted into contiguous token segments (start/end indices) by grouping consecutive states belonging to the same token.

##### Local refinement of token boundaries by grid search

After Viterbi decoding, token boundaries were locally refined using a grid search that optimized alignment between the observed 50-ms z-FR and the corresponding template. For each token segment, we searched over (i) temporal shifts (±0.5 s in 50-ms steps) and (ii) temporal stretch factors (0.4–1.5) applied to the template via linear interpolation. For each candidate shift/stretch, we computed the per-channel Pearson correlation between the stretched template and the observed segment and selected the shift/stretch maximizing the mean correlation across channels. The refined boundaries were used to produce the final token timing estimates and the decoded token sequence. This local boundary refinement procedure follows the approach described in Willett et al. (2021) ^3^.

##### Iterative template updating using decoded segments

To mitigate nonstationarity across sessions, we iteratively refined the templates using decoded segments obtained from forced alignment. Using the refined token boundaries, we extracted the corresponding 50-ms binned z-FR segments and pooled them by token class. For each class, if at least 10 segments were available, we linearly resampled each segment to match the current template length (number of states) and averaged across segments to form an updated class template.

We then repeated the forced-alignment procedure without a Blank state using the updated templates and continued iterating until the frame-normalized Viterbi score (average log-likelihood per frame) no longer improved, with early stopping after three consecutive non-improving iterations. The resulting refined 30-class templates were saved and subsequently used to initialize the final forced-alignment stage that included a Blank state (31-class HMM).

#### Final forced-alignment labeling with a Blank state (session-wise implementation)

Final forced-alignment labeling was performed independently for each closed copy-typing session. This stage used the same HMM structure, emission model, and decoding procedures described above, with the explicit inclusion of a Blank (none) state between successive tokens.

The Blank state was introduced at the final labeling stage for two reasons. First, the online RNN decoder predicted a state space that explicitly included a Blank class, and incorporating a corresponding state in the HMM ensured consistency between the neural decoder outputs and the offline forced-alignment model. Second, explicitly modeling a Blank state was expected to improve segmentation by capturing periods in which the participant was not attempting any finger movement, thereby reducing spurious assignments of movement states during inter-token intervals.

For each session, refined movement templates obtained from the preceding template-refinement stage (30 finger-movement classes) were combined with the original Blank template, yielding a total of 31 templates. These templates were used to build a left-to-right HMM constrained by the ground-truth token sequence for each prompted sentence, with a Blank state explicitly inserted between each pair of adjacent tokens.

For each sentence, emission log-likelihoods were computed under an isotropic Gaussian model (*σ*^2^*I*, *σ*^2^ = 1) by comparing the observed z-FR with the state-wise template means. Viterbi decoding was then performed with temporal window constraints and a terminal constraint enforcing termination in the final Blank state.

Template lengths were adjusted per sentence only when the total number of template states exceeded the number of observed time bins by a predefined margin (*L_µ_*/*T* ≥1.2). When this occurred, the duration of each token template was matched to the observation length by linearly resampling the template along the time axis. Specifically, a global scaling factor *g* was computed from the ratio of the observed sequence length *T* to the total number of template states (*L_µ_*), and each token template was resampled to a new length *L*_out_ using linear interpolation. Blank templates were compressed more aggressively using an additional weighting factor (*w*_blank_ =0.2).

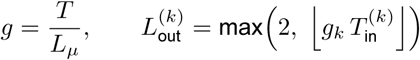

where 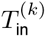 is the original number of states in the template for token *k*, and *g*_k_ = *g* for finger-movement tokens and *g*_k_ = *gw*_blank_ for the Blank state.

This procedure ensured that the emission-mean sequence did not exceed the length of the observed neural data and enabled feasible forced alignment as typing durations shortened across sessions.

Decoded state sequences were converted into token-level segments, and token boundaries were further refined using a local grid search over temporal shifts and linear stretch factors, maximizing similarity between observed neural activity and the corresponding templates.

#### Evaluation of HMM-derived token boundaries using RNN outputs

Because the participant’s intended keystroke timing and token identity during closed-loop copy typing were not directly observable, we evaluated the validity of the HMM-derived token boundaries using the online RNN decoder outputs as an external reference.

##### Character-level accuracy assessment using CER

Character-level accuracy for both methods was quantified using the CER, defined as the normalized number of insertions, deletions, and substitutions required to reconstruct the prompted sentence. CER was computed separately for the character sequences obtained from forced-alignment HMM labeling (Figure 2E) and from the top-1 RNN predictions (Figure 2F).

##### Temporal matching of HMM and RNN token boundaries

To assess temporal consistency between the two methods, we treated the RNN output as a proxy for the participant’s intended token sequence and compared it against the HMM-derived labels on a token-by-token basis. For each sentence, HMM-derived token boundaries were aligned with RNN-derived token boundaries using a greedy, label-matched temporal pairing procedure.

##### Definition of matched tokens

Specifically, an HMM token was considered a matched token if it satisfied two criteria: (i) the token label matched the corresponding RNN output token, and (ii) the HMM-derived token onset preceded the RNN-derived onset by at least 0 s but no more than 1.5 s. The upper bound (1.5 s) was chosen heuristically, informed by the temporal integration of the RNN decoder (300 ms), to retain a sufficient number of matched tokens. Tokens violating this temporal window were excluded as temporal mismatches (Figure 2B, H). Tokens that satisfied the temporal matching criteria were classified as matched tokens, representing neural events consistently identified by both methods.

##### Typing speed and accuracy metrics

Typing speed was quantified as characters per minute (CPM) based on the RNN output during closed-loop copy typing (Figure 2I). For each prompted sentence, CPM was computed by dividing the total number of tokens emitted by the RNN decoder by the total typing duration for that sentence (in minutes), yielding a single CPM estimate per sentence. This metric did not account for the correctness of the decoded characters and therefore served as a measure of typing speed rather than accuracy. For matched tokens only, token-wise typing duration (s) was defined as the difference between the HMM-derived token end time and start time (Supplemental Figure S6A, B). Typing accuracy was assessed separately using the proportion of matched tokens, defined as the number of matched tokens divided by the total number of prompted tokens for each sentence (Figure 2G). The relationship between typing speed and typing accuracy was evaluated using Spearman correlation and linear regression (Figure 2K, L; Supplemental Figure S6C, D). In addition, within-session changes in typing speed across prompted sentences were evaluated using linear regression (Figure 2J).

Only matched tokens were retained for subsequent analyses, as these tokens were most likely to reflect neural activity associated with the participant’s intended keystrokes. The daily proportion of each token was calculated as the number of matched occurrences of that token divided by the total number of matched tokens within the session (Figure 2C).

#### Clipped closed-typing data based on HMM labeling

For each matched token, neural activity was clipped using the HMM-derived start and end times, and analyses were performed on z-FR. FR were computed from ncTX using a Gaussian kernel (*σ* = 40 ms) and z-scored using the session-specific rest mean and standard deviation (see “Preprocessing of typing-session neural activity”).

Because HMM-derived token durations varied across occurrences, clipped z-FR segments had variable lengths even within the same token class. To enable within-token alignment, we grouped clipped segments by token identity and constructed a trial × time × channel data structure for each token by zero-padding each trial to the maximum segment length observed for that token within the session. Because z-FR is centered at zero by construction, zero-padded samples correspond to baseline activity rather than artificial suppression of firing.

##### Refinement of clipped z-FR within each token

To refine grouped clipped signals within each token (trial × time × channel data), we applied a piecewise time-warping procedure using the `PiecewiseWarping` model (see “Template construction for forced-alignment labeling using an HMM”). Time-warping was applied separately within each token class to align waveforms across trials after zero-padding.

Following time-warping, we performed a dedicated refinement step to remove padding-dominated regions. For each token, we computed trial-averaged responses (PSTHs) from the time-warped z-FR and visually inspected the averaged waveforms across channels to identify portions of the aligned response that reflected genuine neural activity rather than padded samples (Supplemental Figure S8A). Based on this inspection, we defined a token-specific valid time range and trimmed the time-warped z-FR to that range, yielding a refined z-FR dataset for each token. Subsequent analyses of across-session changes used these refined, clipped z-FR datasets (Figure 5, 6, 7).

##### Baseline z-FR construction

To construct baseline activity matched to each sentence, we extracted z-FR from the preparation period preceding copy-typing. For each sentence, the preparation segment was used as the baseline reference for all matched tokens in that sentence. To avoid early cue-evoked transients, baseline z-FR was extracted from a fixed window 1.01–2.00 s after preparation cue onset (time bins 101–200; 10 ms bins) (Supplemental Figure S8B).

Because the baseline window was fixed in duration (100 data points) across all trials, baseline segments had identical lengths within each token class. As a result, zero-padding, time-warping, and visual trimming steps were not required for baseline z-FR construction. These baseline, refined z-FR data were used in paired t-tests for spatiotemporal cluster analysis (Figure 5F, G).

#### Neural representations and dynamics during closed copy-typing

All subsequent analyses of closed copy-typing data were performed using the refined, token-aligned z-FR representations described above. These analyses examined how neural dynamics, representational geometry, population structure, and inter-channel relationships evolved over the course of closed copy-typing sessions.

#### Population representational structure across sessions evaluated by Mahalanobis distance

Closed copy-typing sessions were analyzed across seven recording days (Day 45, 85, 86, 92, 93, 106, and 107). By design, the copy-typing task involved 30 distinct finger-movement tokens; however, not all tokens were successfully segmented and matched in every session (Figure 2C). Accordingly, all analyses were restricted to tokens that were actually typed within each session. To ensure sufficient sampling, tokens with insufficient occurrences were excluded; any token with fewer than 10 valid trials within a session was omitted from subsequent analyses. Unless otherwise specified, all analyses were performed separately for right 6d and left 6d.

##### Session-wise construction of population feature vectors

For each session and token, we assembled a trial-by-time-by-channel array from the refined z-FR. Although neural activity was temporally aligned within each token segment, token durations were not aligned across tokens. To focus on population-level representational structure rather than temporal dynamics, neural activity within each token was averaged across time, yielding a single population feature vector per trial.

##### Quantifying token separability using Mahalanobis distance

To quantify the separability of token-evoked population representations, we computed pairwise normalized Mahalanobis distances between tokens within each session (Figure 5A; see “Mahalanobis distance” for methodological details). Because the Finger Movement task demonstrated that right 6d strongly encodes right-versus-left hand information, resulting in a clear hierarchical structure in dendrogram form (Figure 3F), we illustrated right 6d dendrograms to assess whether population representational structure was preserved across sessions (Figure 5B, Supplemental Figure S9).

##### Stability of population representational geometry across sessions

To quantify changes in population representational structure across sessions, we plotted session-to-session changes in normalized Mahalanobis distances (Figure 5C, left axis). Group-wise differences across sessions were evaluated using the Friedman test. To assess the stability of representational geometry across adjacent sessions, we computed Spearman correlations between normalized Mahalanobis distance matrices from consecutive sessions (e.g., Day 45 vs. Day 85, Day 85 vs. Day 86; Figure 5C, right axis).

##### Relationship between Mahalanobis separability and decoding performance

Because we previously observed a strong positive relationship between Mahalanobis separability and SVM decoding performance in the Finger Movement task (Figure 3H), we additionally trained and evaluated across-session SVM classifiers using the refined z-FR features (Figure 5D).

##### Inter-areal comparison of representational geometry between right and left 6d

To examine how representational geometry in right 6d and left 6d related across sessions, we compared their session-wise pairwise-distance structures using two complementary approaches. First, for each token pair, we quantified inter-area divergence by computing the absolute difference between right 6d and left 6d normalized Mahalanobis distances; differences across sessions were evaluated using the KW test. Second, within each session, we computed the Spearman correlation between right 6d and left 6d normalized Mahalanobis distance matrices, providing a measure of similarity in population representational structure between the two areas (Figure 5E).

#### Spatiotemporal cluster analysis of refined z-FR during closed copy-typing

We quantified session-wise changes in refined z-FR during closed copy-typing by comparing baseline and typing phases at the channel level using spatiotemporal cluster analysis implemented by FieldTrip (see “Cluster-based spatiotemporal permutation test” for methodological details). Within each segmented token, refined z-FR was averaged across time to yield one population vector per trial (trial × channel). Tokens with fewer than 10 valid trials within a session were excluded from subsequent analyses. Paired t-tests were performed across trials for each channel, and multiple comparisons were controlled using spatiotemporal cluster-based correction (Supplemental Figure S10).

Significant positive t-values were classified as FR activation, and significant negative t-values as FR suppression. Finger-movement tokens were grouped into right-hand and left-hand finger sets. For each session and for right 6d and left 6d, we plotted the normalized mean t-value across sessions, computed separately for activation and suppression (Figure 5F,G). Differences across sessions were assessed using the KW tests.

#### Functional connectivity analyses

We quantified functional connectivity within and between right 6d and left 6d using a predictive modeling framework reported by Perich et al., 2018 ^19^. Refined z-FR trajectories within each token were aligned by time warping and had matched durations within a session, but varied across tokens and across sessions. Therefore, for each finger-movement token, we computed PSTHs as the trial-averaged neural activity over time, requiring at least 10 trials per token (tokens with fewer trials were excluded). PSTHs were then linearly stretched to a fixed template length using one-dimensional linear interpolation applied independently to each channel (typically 20 bins for dimensionality estimation and 100 bins for connectivity analyses), yielding a set of token-wise PSTHs with equal time length.

Dimensionality estimation used the shorter template (20 bins) because the goal was to compare task-averaged population structure, whereas generalized linear model (GLM) and canonical correlation analysis (CCA) used the longer template (100 bins) to stabilize prediction performance.

#### Noise-corrected dimensionality estimation

To determine how many PCA dimensions reflected signal rather than trial-to-trial noise, we estimated a noise floor using surrogate data ^19^. For each movement condition, trials were randomly permuted and split into pairs; for each pair, we computed two PSTHs and defined a noise sample as their difference, capturing trial-to-trial surrogate noise. This procedure was repeated 1,000 times to generate a distribution of noise PCA eigenvalues for each dimension. For each dimension, the 99th percentile of surrogate eigenvalues was taken as the maximal variance explainable by noise. Signal eigenvalues were estimated by subtracting this noise upper bound from the real-data PCA eigenvalues, with negative values set to zero. The number of significant principal components (PCs) was defined as the smallest number of PCs explaining 95% of the total signal variance.

We computed this noise-corrected dimensionality separately for right- and left-hand movement conditions in right and left 6d, with signal-related component counts calculated within each session (Figure 6A). Differences in the number of signal-related PCs between right and left 6d were assessed using the Wilcoxon signed-rank test.

#### Intra-array connectivity using predictive GLMs

To restrict connectivity analyses to class-selective channels, we performed the KW tests combined with FDR correction (see “Selection of class-selective channels”). For within-day connectivity, channel selection was performed within that session day. For within-day analyses, channel selection was performed separately for each session. For cross-day analyses, only channels that were significant across all included sessions were used. Analyses were further restricted to finger-movement tokens common across all session days.

##### Within-day intra-array connectivity

For each session and area (right 6d, left 6d), we quantified intra-array connectivity by predicting each target channel from the population activity of the remaining channels. PSTHs were concatenated across tokens to form the predictor matrix. For each target channel, that channel was excluded from the predictor set. Based on the dimensionality estimation results (Figure 6A), we retained the top 10 PCs as signal components and fit a linear Gaussian GLM (MATLAB `fitglm`) (Supplemental Figure S11). Prediction performance was evaluated using 10-fold cross-validation, with PCA recomputed within each fold to prevent information leakage. Connectivity strength was quantified as the distribution and mean of cross-validated normalized root mean squared error (NRMSE) values across target channels.

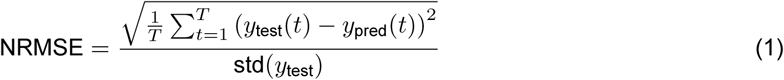

An NRMSE value of 0 indicates perfect prediction (i.e., identical predicted and true signals), with larger values indicating poorer prediction performance. Differences across sessions were assessed using the KW tests (Figure 6B).

##### Cross-day intra-area connectivity (forward vs backward prediction)

To assess the stability of intra-area coupling across sessions, we trained the GLM on one day and evaluated prediction on another day. For each pair of days, two prediction directions were evaluated:

- Forward prediction: training on the earlier day and testing on the later day
- Backward prediction: training on the later day and testing on the earlier day

For each day pair and target channel, the GLM was trained in a common PCA space derived from concatenated data across all days (computed after excluding the target channel). Both training and test data were projected into this space using the top 10 PCs, and cross-day NRMSE values were computed. Forward and backward prediction performance was compared. Based on better performance observed for backward prediction (Figure 6C), subsequent area comparisons focused on backward prediction to assess right- and left-hand group performance within right and left 6d (Figure 6D). Differences between paired group prediction performance were evaluated using the Wilcoxon signed-rank test.

#### Inter-area connectivity using CCA in PCA space

We performed CCA between right 6d and left 6d activity in PCA space. For each condition (right-hand or left-hand movements), we constructed concatenated PSTH matrices separately for right and left 6d using the across-day common token set and channels significant across all sessions. Within each session, PCA was computed separately for right and left 6d matrices, and the top 10 PCs were retained. Canonical correlations were then computed between the right and left 6d matrices (Figure 6E).

Because higher-order canonical dimensions were noisier, CCA results were summarized using the top eight canonical components, chosen as the smallest set of components that captured close to 90% of the total shared variance in the data (Supplemental Figure S12). Confidence intervals on the mean canonical correlation were estimated using bootstrap resampling, and differences across sessions were assessed using a Friedman test. Day-wise mean ranks were visualized for the summary.

#### Inter-area connectivity using predictive GLMs

We quantified directional inter-area connectivity by predicting channel activity in one 6d from population activity in the other 6d within each session. Two predictive models were fit:

- *R* → *L*: predictors from right 6d, targets in left 6d
- *L* → *R*: predictors from left 6d, targets in right 6d

For each target channel, predictors were reduced to the top 10 PCs computed from the predictor population, and a linear GLM was trained. Prediction performance was assessed using 10-fold cross-validation, yielding a distribution of NRMSE values across target channels. Differences across sessions were evaluated using the KW tests (Figure 6F).

#### dPCA analysis of closed-loop finger typing dynamics

Specific calculation procedures and analysis settings followed those described for the open-loop finger movement task (see “dPCA of the Finger Movement task”). Because not all tokens were observed in every closed-loop session(Figure 2C), we restricted analyses to tokens for which (i) all three gestures (Up/Down/In) were present and (ii) the corresponding finger was observed for both hands. Under these constraints, the only fingers consistently available across all closed-loop sessions were Thumb and Middle; therefore, the Finger marginalization in the closed-loop analyses included only these two fingers (in contrast to the open-loop task, which included five fingers).

##### Equalizing token duration across trials

Token durations derived from the clipped, refined z-FR data varied across tokens. To equalize durations prior to dPCA, each trial trajectory was linearly resampled to a fixed template length of 100 bins using one-dimensional linear interpolation. This produced trial matrices of shape [100 × channels]. The same template length was used for all sessions. We then performed dPCA on condition-averaged PSTHs and projected the PSTHs onto the corresponding dPCA axes. For visualization and subsequent analyses, we focused on the top two components (dPC1–2) for each marginalization. Session-wise trajectories in the dPC1–2 plane were visualized as time-ordered curves for each marginalization (Figure 7A).

##### Latent trajectory length (“magnitude”) in dPC1–2

To quantify the scale of latent dynamics, we computed the trajectory length in the dPC1–2 plane for each condition as the sum of Euclidean step-wise lengths across consecutive time points; this metric is hereafter referred to as the trajectory “magnitude”.

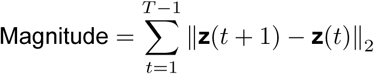

where **z**(*t*) ∈ ℝ^2^ denotes the dPC1–2 latent state at time *t*.

We computed this length for each token and, within each session, averaged lengths across subgroups within each marginalization (Hand, Finger, and Gesture). To test for session-wise changes in trajectory length, we used the KW test across sessions (Figure 7B).

##### Relationship to typing performance

To relate latent magnitude to behavioral and decoder performance, we compared session-wise mean trajectory length to (i) typing speed (CPM) and (ii) decoder performance (RNN CER). We computed Pearson correlations between magnitude and each performance metric and visualized linear fits for interpretability (Figure 7C).

#### Across-session alignment of latent dynamics using CCA

We tested whether session-specific latent trajectories could be linearly aligned using canonical correlation analysis (CCA). If session-to-session changes can be captured by CCA alignment, then aligning latent trajectories from two sessions into a shared subspace should reduce systematic relationships between inter-session interval and differences in latent trajectory magnitude.

For the seven closed-loop sessions, we enumerated all unique pairs of sessions. For each pair, we computed the inter-day interval as the absolute difference in days between the two sessions. All alignment and comparison analyses were performed separately for right 6d and left 6d areas and for each marginalization. Our alignment procedure followed the approach described by Gallego et al., 2020 ^20^.

##### CCA-based alignment of dPC trajectories

For each session pair, token, and marginalization, we extracted the session-specific dPC1–2 trajectories,

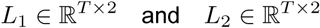

where *T* denotes the number of time points.

Each trajectory was mean-centered across time, and CCA was performed using MATLAB’s canoncorr function:

- *X* = *L*_1_ − mean(*L*_1_)
- *Y* = *L*_2_ − mean(*L*_2_)
- [*A, B, r*] = canoncorr(*X, Y*)

The aligned latent trajectories were then obtained by projecting the original trajectories into the CCA space:

- *U* = *XA*
- *V* = *YB*

These aligned trajectories (*U, V*) were used as the aligned latent dynamics for all subsequent analyses.

##### Differences in latent trajectory magnitude across sessions (unaligned vs aligned)

For each session pair and token, we computed the latent trajectory magnitude separately for each session using the unaligned dPC1–2 trajectories and again after CCA alignment. We then computed the absolute difference in trajectory magnitude between the two sessions.

To assess whether differences in latent trajectory magnitude increased with inter-day interval, we related the absolute magnitude difference to the inter-day interval using Spearman’s rank correlation coefficient. These relationships were visualized using scatter plots with linear fits for unaligned (blue) and aligned (red) trajectories (Figure 7D, left panels).

In addition, we compared the distributions of unaligned and aligned magnitude differences using histograms and assessed statistical differences using a paired Wilcoxon signed-rank test, reflecting the paired structure of unaligned and aligned values for each session pair (Figure 7D, right panels).

#### Statistical analysis

Statistical analyses were performed using nonparametric tests unless otherwise noted. For paired comparisons across multiple conditions, the Friedman test was used. For paired comparisons between two conditions, the Wilcoxon signed-rank test was applied. For unpaired comparisons across multiple groups, the KW tests were applied. Spatiotemporal cluster-based analyses were conducted using paired t-tests at the sample level, with cluster-level inference to control for multiple comparisons. Associations between variables were assessed using Spearman’s rank correlation coefficient. To correct for multiple comparisons, spatiotemporal cluster-based correction and FDR correction (*α* = 0.05) were applied as appropriate. Statistical significance was defined as *p<* 0.05. For clarity and consistency, all p-values smaller than 0.001 are reported as *p<* 0.001.

