## Supplementary Information for "Learning-related population dynamics in right and left dorsal premotor cortex during typing skill acquisition"

**Supplemental Figure**

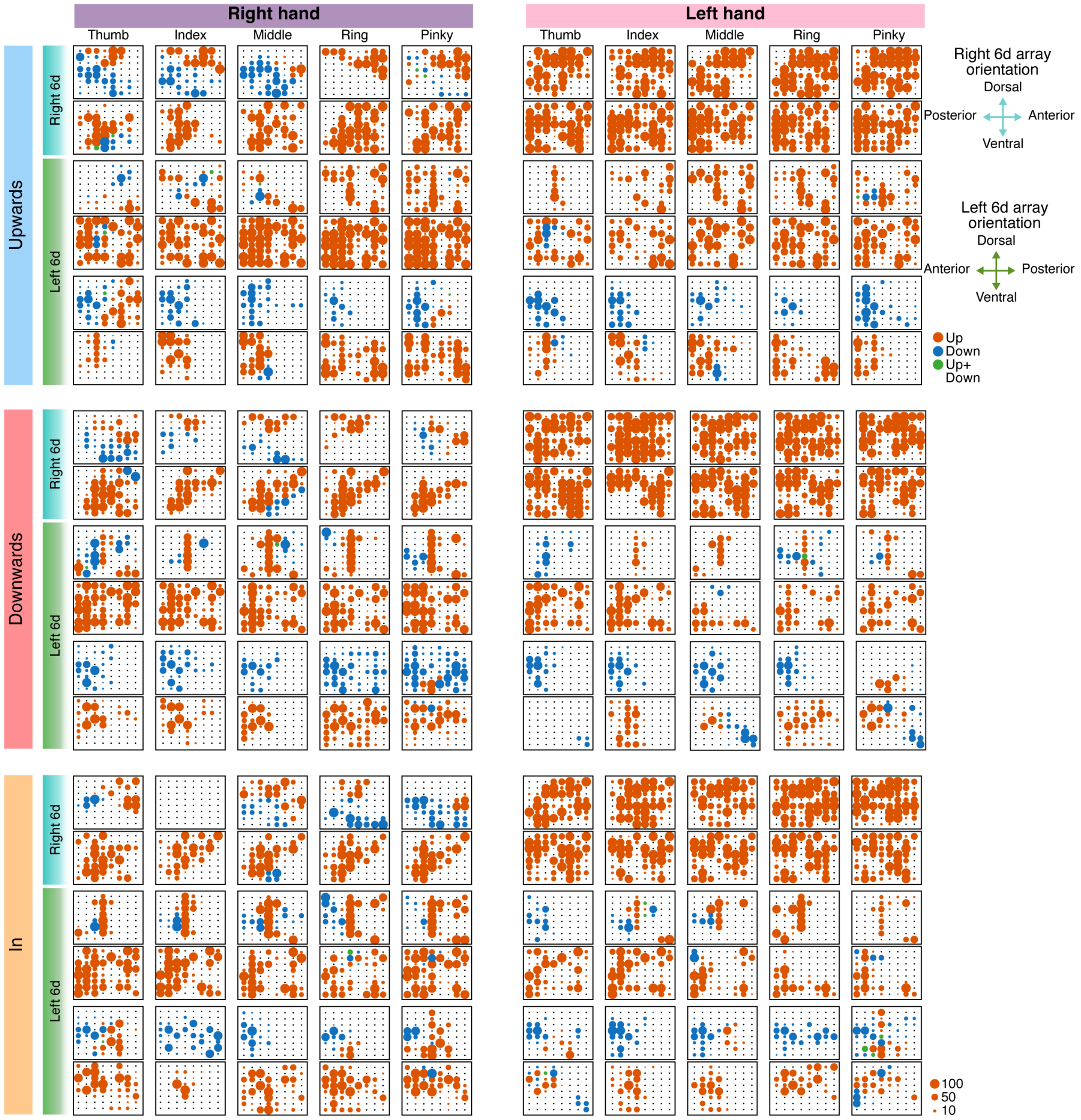


**Supplemental Figure S1: Spatiotemporal cluster analysis across all 30 finger movements.**
Spatiotemporal cluster analysis results for all 30 finger movement conditions (five fingers × three movement directions for each hand). Neural activity from 0–1 s after the Go was analyzed. Each panel shows the spatial distribution of electrodes in dorsal premotor cortex (Brodmann 6d) exhibiting a significant increase (red), decrease (blue), or both (green) relative to baseline.
The size of each circle reflects the number of time bins (out of 100 bins spanning 0–1 s) in which the electrode was part of a significant cluster, providing a measure of the temporal extent of modulation. Panels are organized by movement direction (Up, Down, In), hand (right vs. left), finger (thumb to pinky), and hemisphere (right vs. left 6d).
Notably, during left-hand movements, modulation in right 6d was predominantly characterized by increases in activity, whereas during right-hand movements, both increases and decreases were observed in right 6d.
Anatomical orientations of the NeuroPort electrodes for right and left 6d are indicated.
All data shown in this figure were obtained during the Finger Movement task on trial day 43.

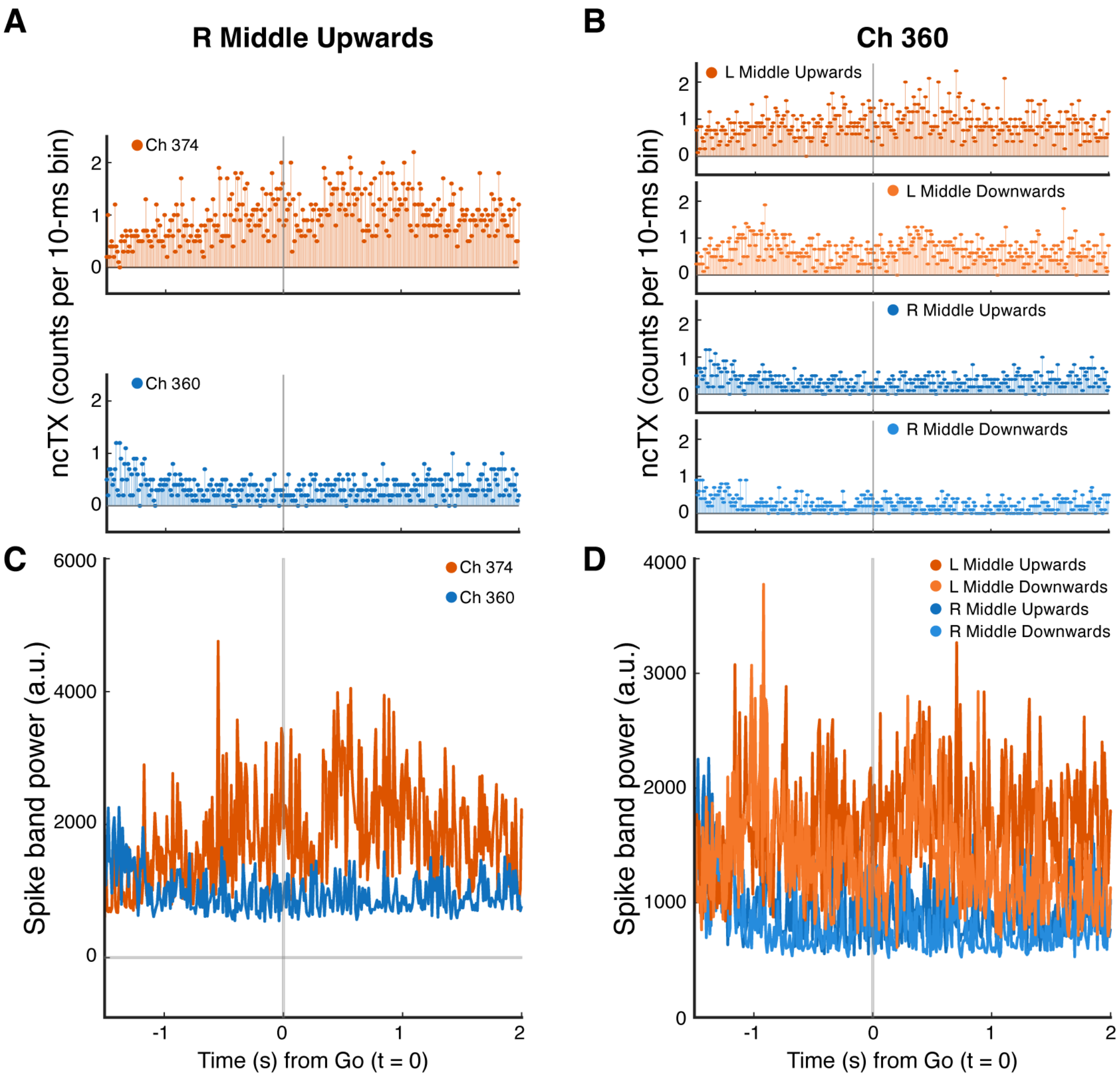


**Supplemental Figure S2: Example trial-averaged neural responses represented using ncTX and spike-band power**.

Panels A and B show trial-averaged ncTX for the same example channels and movement conditions shown in Figure 1E. Panels C and D show the corresponding trial-averaged spike-band power signals. The example channels illustrate both increases and decreases in neural activity associated with finger movements. Compared with threshold crossing rate representations (Figure 1E), ncTX and spike-band power signals exhibit greater bin-to-bin variability, making temporal response profiles more difficult to visualize. For this reason, threshold crossing rates were used in the main figures and are referred to throughout the manuscript as firing rates (FR).

All data shown in this figure were obtained during the Finger Movement task on trial day 43.

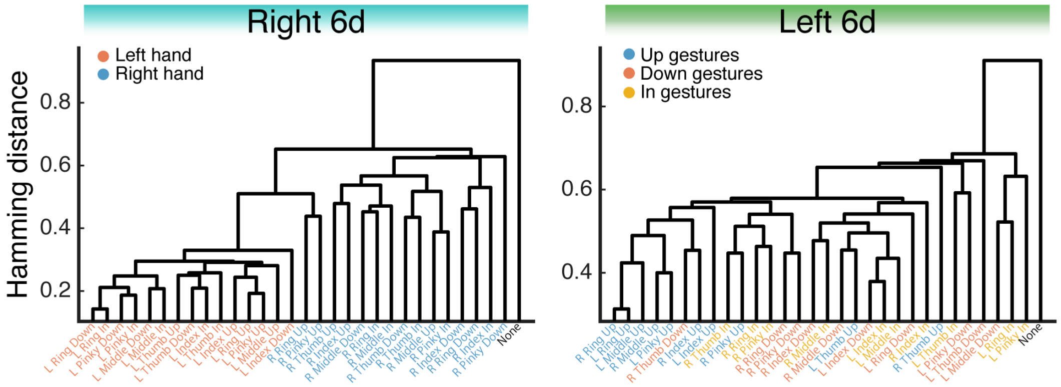


**Supplemental Figure S3: Dendrograms constructed using average linkage**.
Dendrograms were generated using the same masked Hamming distance matrices and task conditions as in Figure 1H, but with average linkage instead of Ward linkage. Leaf labels are color-coded according to hand (right vs. left) for right 6d (left) and movement direction (up, down, in) for left 6d (right). No substantive differences in the overall clustering patterns were apparent compared to Figure 1H.

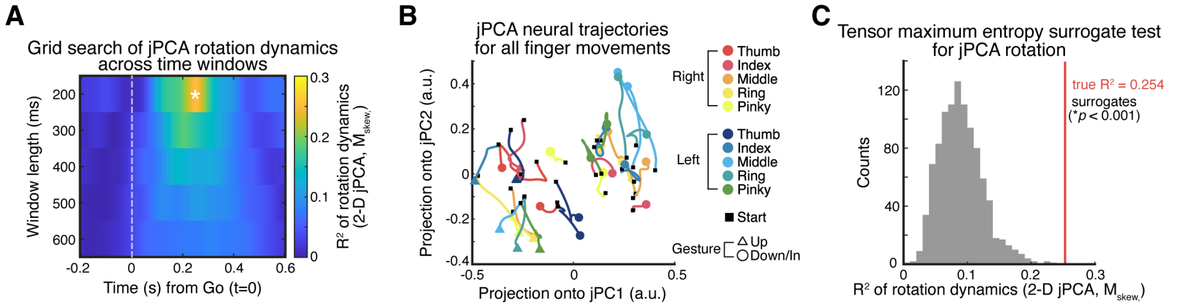


**Supplemental Figure S4: Low-dimensional rotational neural representations of finger movements in bilateral 6d**.
We examined population-level dynamics using jPCA applied to z-scored firing rates (z-FR). Because jPCA identifies rotational structure shared across conditions, it was applied to z-FR from all arrays and all 30 finger movements. Rotational strength was quantified using the coefficient of determination ($R^2$) for the skew-symmetric component in the jPC1–2 plane. A grid search across temporal windows revealed a brief peak in rotational dynamics when using a 200-ms window centered at 0.25 s after the Go (A). Using these parameters, condition trajectories exhibited a clear rotational pattern, with Up gestures clustering in the third quadrant and Down/In gestures in the first (B). Tensor maximum-entropy surrogate testing confirmed that the observed rotational dynamics were significantly stronger than expected by chance (C).
(A) Grid search of jPCA rotational dynamics across window lengths (200–600 ms) and analysis times (−0.2 to 0.6 s relative to the Go cue). A transient increase in rotational dynamics was observed at 0.25 s using a 200-ms window (white asterisk). jPCA was computed using z-scored FR from all arrays and all 30 finger movements.
(B) jPCA trajectories computed using the 200-ms window centered at 0.25 s (panel A, white asterisk). Upward finger movements were predominantly localized in the third quadrant of the jPCA plane for both hands.
(C) Tensor maximum-entropy surrogate test evaluating the significance of rotational dynamics. The observed $R^2$ value for the 0.25 s / 200-ms window exceeded the upper 0.1% of the surrogate distribution, corresponding to $p < 0.001$.

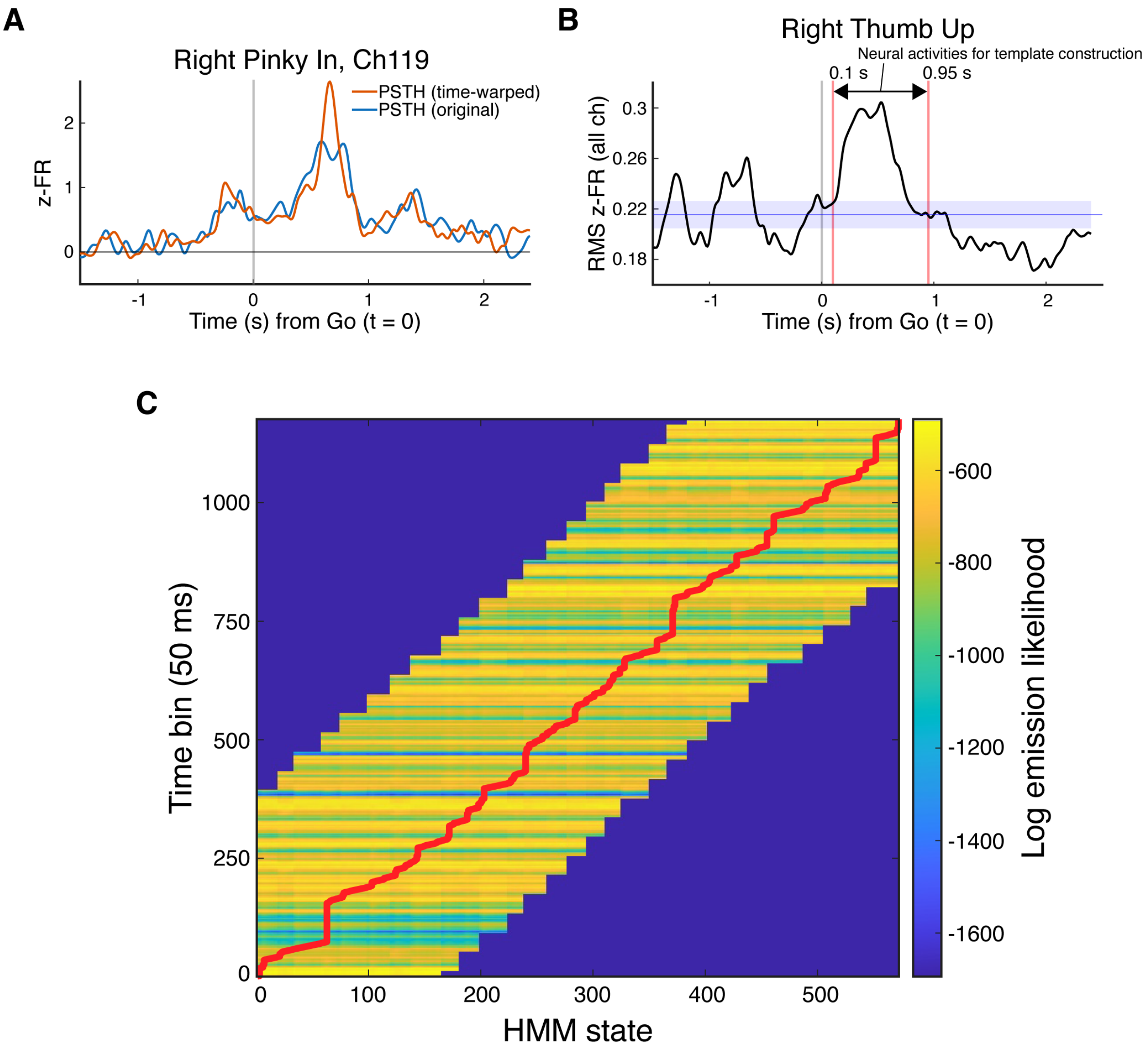


**Supplemental Figure S5: Construction of HMM templates and state inference**.

(A) Example PSTHs from channel 119 during Right Pinky In movements, shown before (blue) and after (red) time-warping. Time-warping aligns movement-related neural activity across trials, resulting in a clearer z-FR profile.

(B) Root-mean-square (RMS) z-scored FR across all channels for Right Thumb Up movements after time-warping. The mean baseline activity during the pre-Go period is shown in blue, with the shaded region indicating ±0.5 SD. Based on visual inspection, the end of movement-related activity was defined as the time point at which RMS activity returned to and remained within the baseline range (mean ± 0.5 SD) for 200–300 ms. Neural activity between 0.1 s and 0.95 s after the Go was used to construct the HMM emission templates.

(C) Log emission likelihood matrix for an example sentence. The x-axis denotes HMM states constructed by concatenating time points from finger movement templates, and the y-axis denotes time bins of the observed neural activity (50 ms bins). Color indicates the log emission likelihood of the observed neural activity given each state. The most likely state sequence inferred by the Viterbi algorithm is shown as a red line. To constrain the search space, allowable states for each token were restricted to a broad temporal window centered on the token's expected relative position within the sentence (±30% of the sentence duration), resulting in masked regions with log likelihood set to $-\infty$.

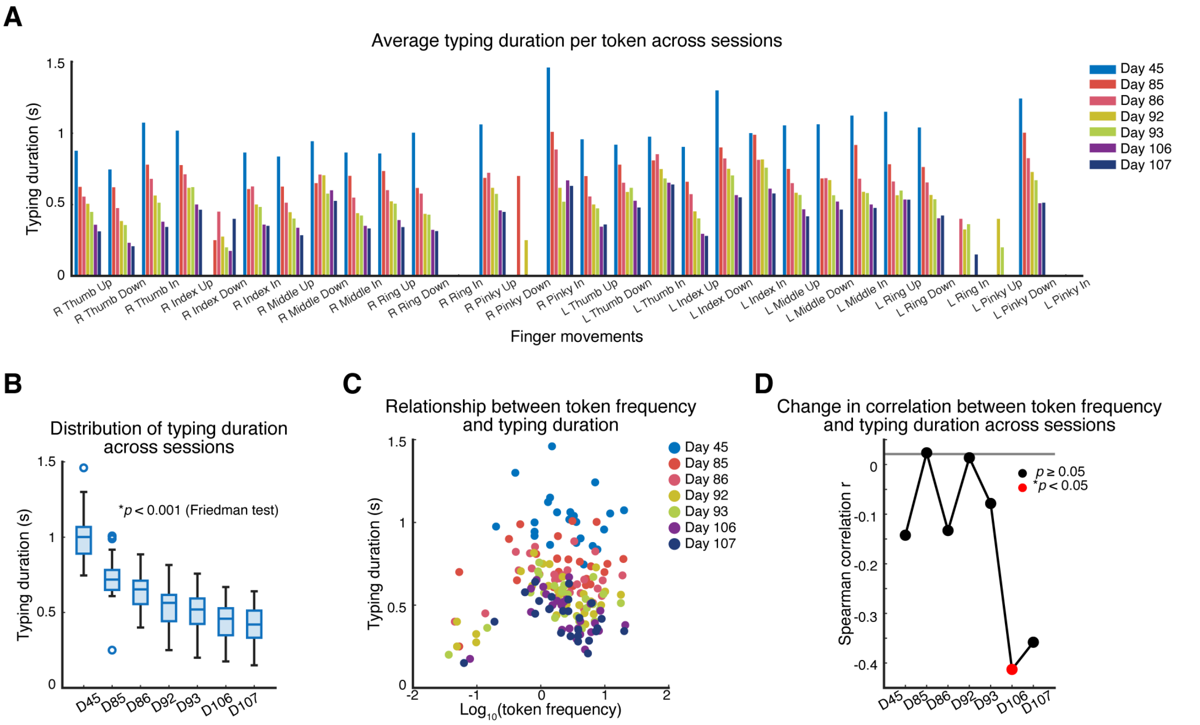


**Supplemental Figure S6: Token frequency does not systematically explain typing speed during closed-loop finger typing**.

(A) Distribution of typing durations for each finger-movement token across closed-loop typing sessions, computed as the interval between HMM forced-alignment onset and offset times. Typing duration tended to decrease across days for nearly all finger movements.

(B) Session-wise changes in typing duration (median ± IQR). Typing durations showed significant modulation across sessions (Friedman test, $p < 0.001$), with a general decreasing trend consistent with the CPM increase shown in Figure 2I.

(C) Scatter plot showing the relationship between token frequency ($\log_{10}$-transformed) and mean typing duration across sessions.

(D) Session-wise Spearman correlations between token frequency and typing duration. Only Day 106 exhibited a significant negative correlation, whereas no significant relationships were observed on other days.

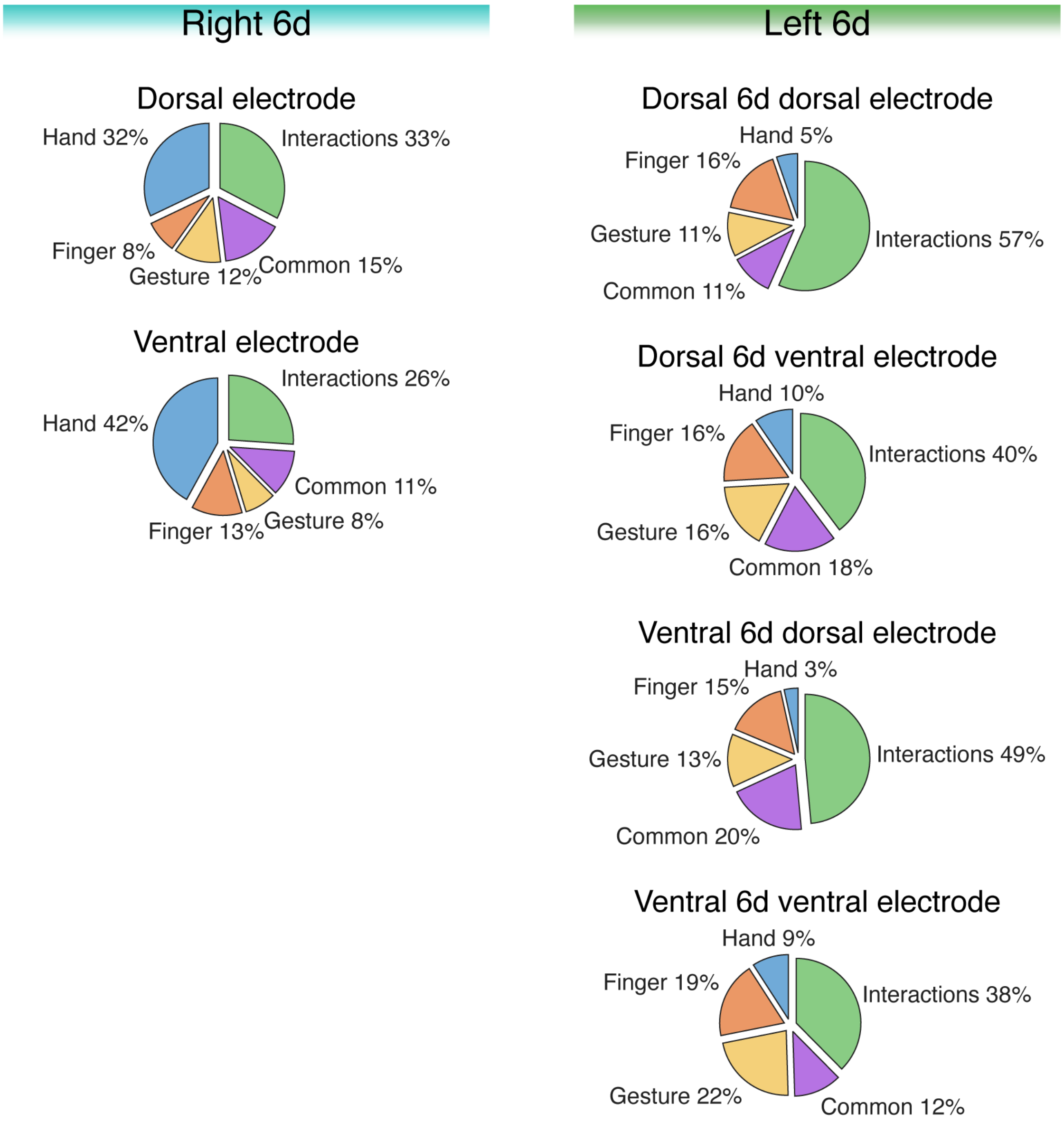


**Supplemental Figure S7: Consistency of demixed population structure across individual premotor electrode arrays**.
dPCA was performed separately for each implanted electrode array in right and left 6d. Pie charts show the proportion of variance explained by Hand, Finger, Gesture, Common, and Interactions for each electrode array. In right 6d (two electrode arrays), Hand marginalizations accounted for a larger fraction of explained variance across both right electrode arrays. In left 6d (four electrode arrays), Finger and Gesture marginalizations were consistently more prominent. These array-specific analyses recapitulated the patterns observed when arrays were pooled within each right or left 6d, indicating that the reported differences in Figure 4C and D were robust across individual electrode arrays.

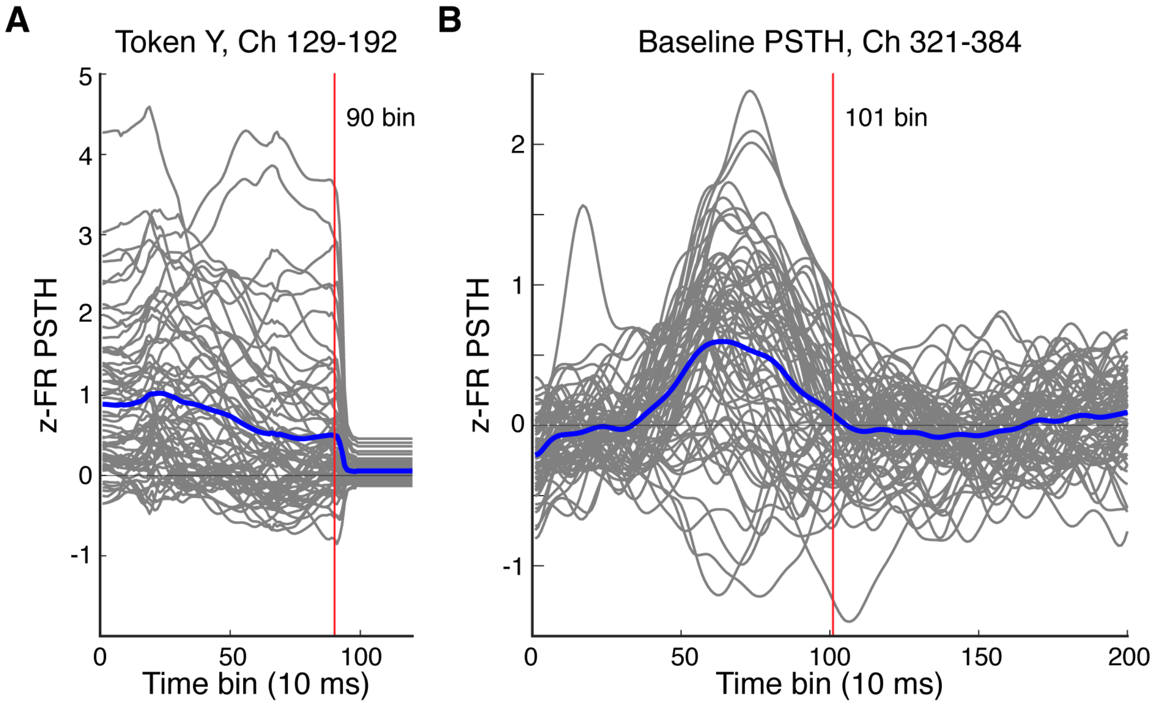


**Supplemental Figure S8: Refinement of HMM-labeled neural data through time-warping**.
(A) Example z-FR activity for Token Y extracted from HMM-labeled data on Day 45. Neural activity was zero-padded to equalize segment length across trials prior to time-warping. PSTHs from channels 129–192 are shown (gray), with the mean PSTH across these channels shown in blue. Because the latter portion of each segment corresponds to zero padding, only the first 90 time bins (red line; 10 ms per bin) were treated as valid data and used for analysis.
(B) Time-warped baseline neural activity from Day 45 for channels 321–384. PSTHs from individual channels are shown in gray, with the mean across channels shown in blue. Transient task-evoked activity is evident during 50–100 bins, indicating that this interval is not suitable for baseline estimation. Accordingly, time bins 101–200 (1010–2000 ms) were used as baseline activity.

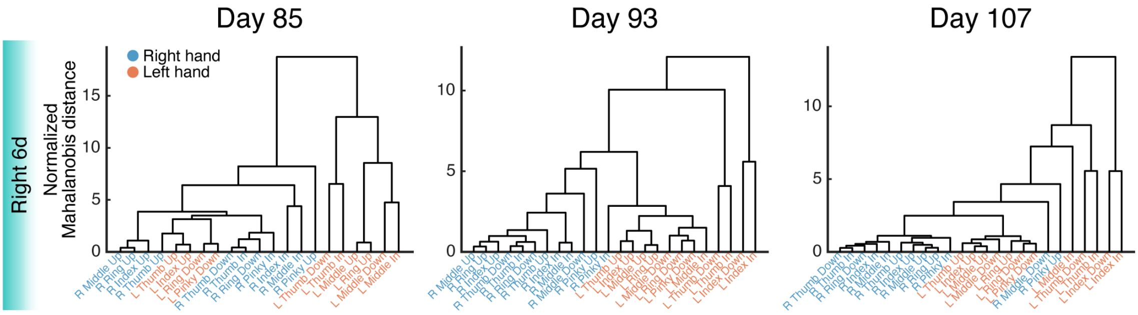


**Supplemental Figure S9: Session-by-session hierarchical clustering of right 6d activity**.
Hierarchical clustering of normalized Mahalanobis distance matrices for right 6d on Days 85, 93, and 107, shown using the same analysis and visualization procedures as in Figure 5B, which displays representative sessions. Leaf labels denote individual finger movements and are color-coded by hand (right hand, blue; left hand, red).

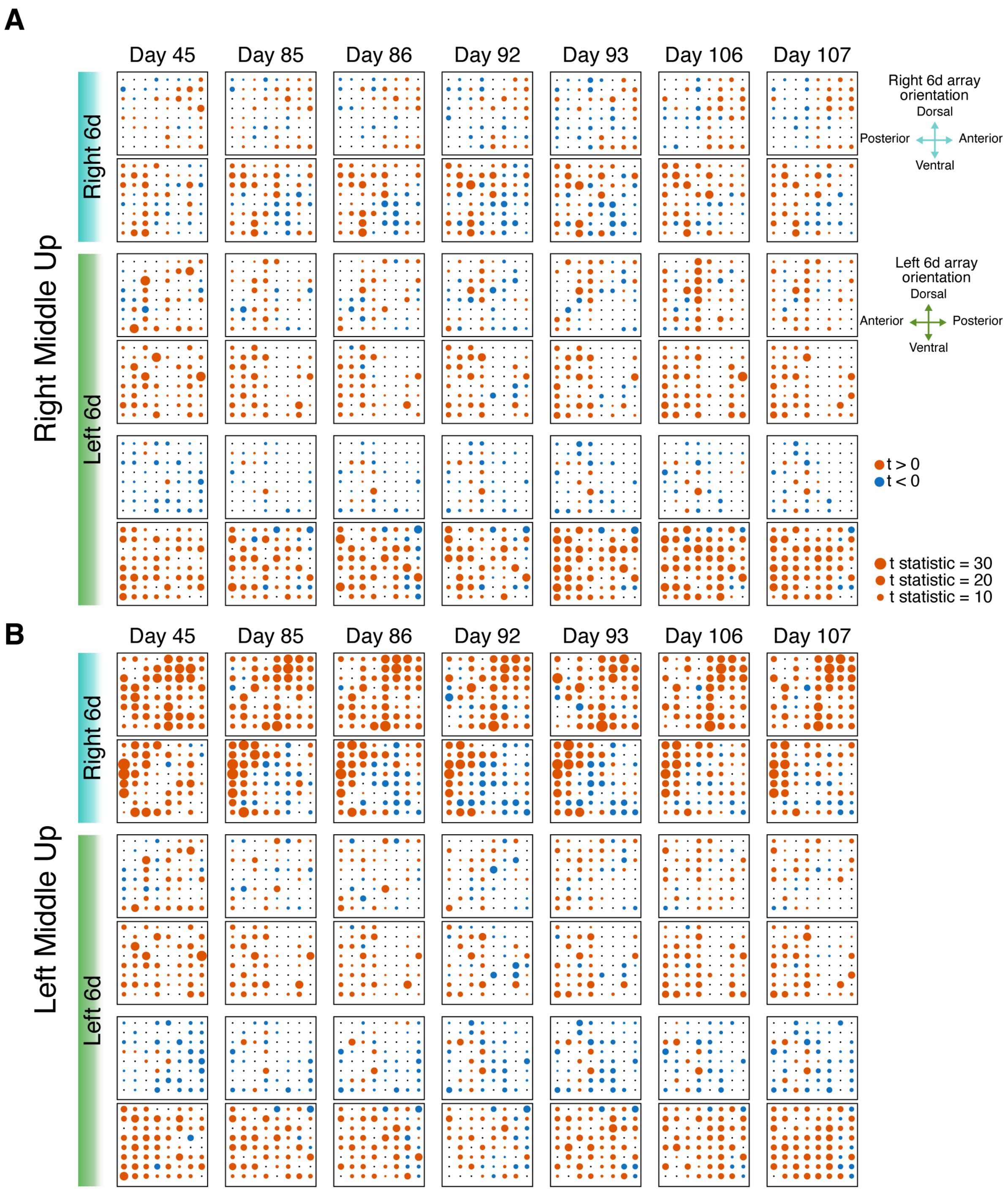


**Supplemental Figure S10: Spatiotemporal cluster analysis of middle finger movements across sessions**.

(A) Spatiotemporal cluster analysis results for Right Middle Up movements across sessions (Days 45, 85, 86, 92, 93, 106, and 107).

(B) Same analysis for Left Middle Up movements.

For each session, refined z-FR extracted using HMM-based labeling was compared against the corresponding baseline activity using a spatiotemporal cluster analysis. The resulting t-statistics are shown for electrodes belonging to significant spatiotemporal clusters in the right and left 6d arrays. Circle size indicates the magnitude of the t-statistic. Red and blue colors denote positive and negative t-values, respectively.

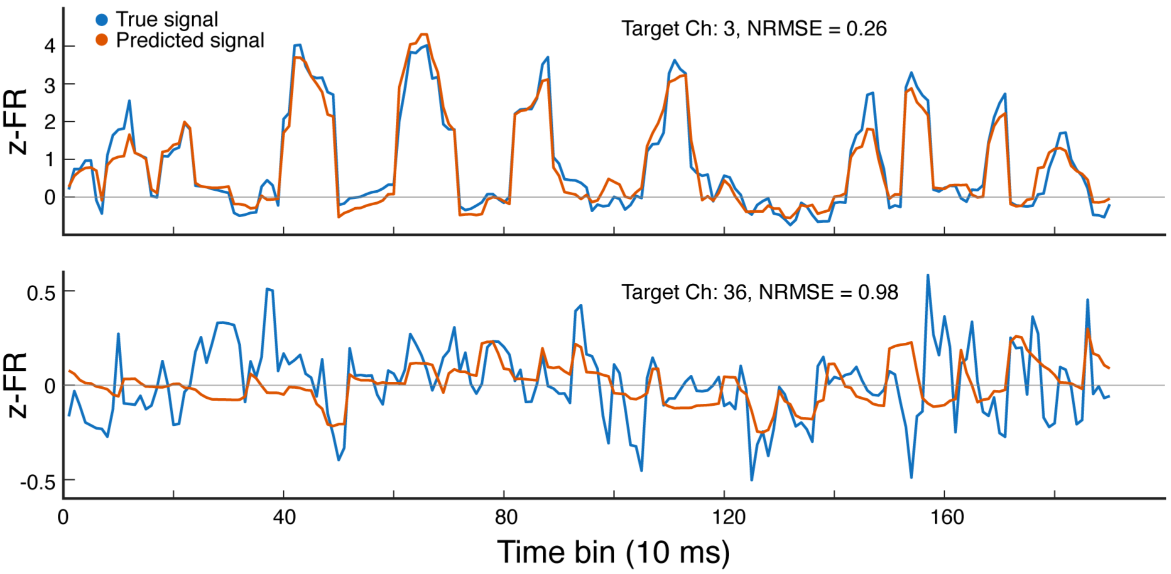


**Supplemental Figure S11: Example GLM prediction of single-channel activity from within-area population signals, illustrating better and worse cases**.

The blue trace shows the true z-FR signal of an example target channel from held-out test data, and the red trace shows the corresponding GLM prediction based on the activity of the remaining channels. The model was trained using the top 10 principal components of population activity. Prediction performance is summarized using the normalized root mean squared error (NRMSE). The better-performing example is shown at the top, and the worse-performing example is shown at the bottom.

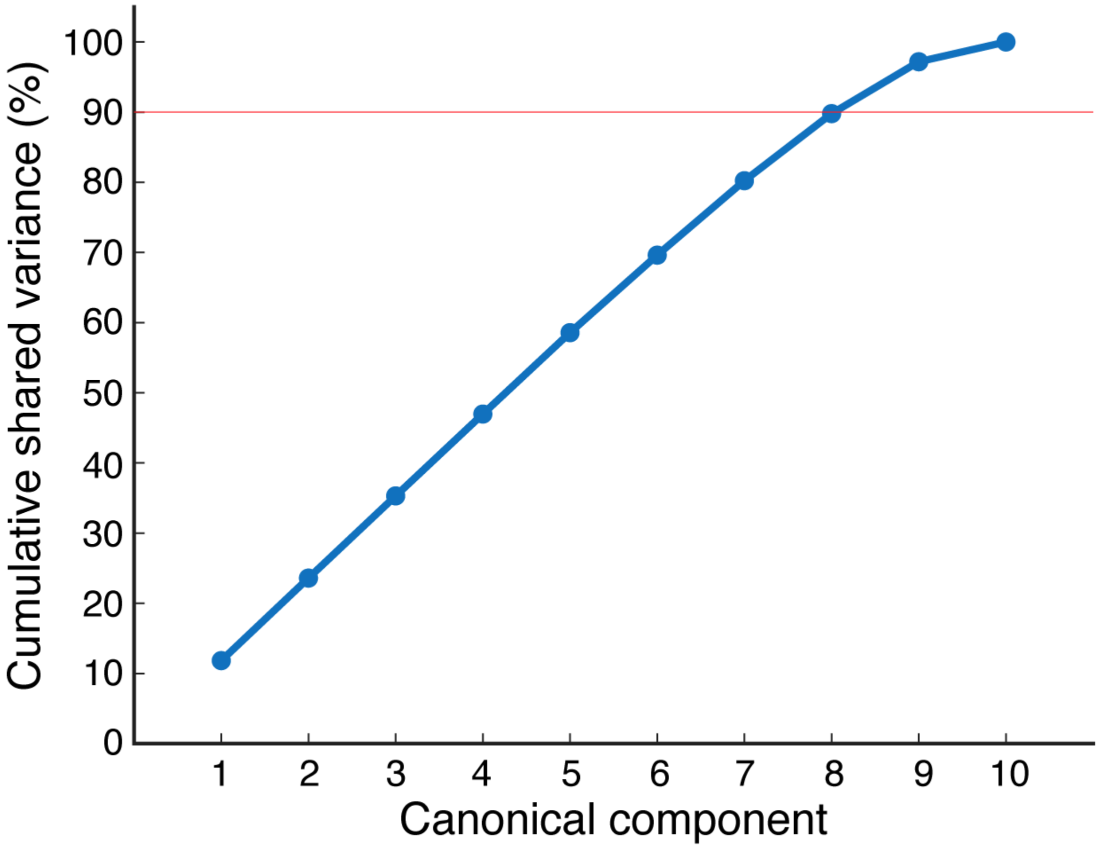


**Supplemental Figure S12: Cumulative shared variance explained by canonical components in CCA**. The cumulative proportion of shared variance explained by successive canonical components is shown. The first eight canonical components together account for approximately 90% of the total shared variance (red horizontal line), motivating the use of these components for Figure 6E.

### Supplemental Table

Supplemental Table S1: T18 Session Data

| Trial Day | Tasks | Number of Sentences |
|---|---|---|
| 43 | 30 Finger Movements Task | 0 +<br>10 repetitions of each isolated finger movement condition |
| 45 | Closed Sentence Copy Typing | 43 |
| 85 | Closed Sentence Copy Typing | 82 |
| 86 | Closed Sentence Copy Typing | 90 |
| 92 | Closed Sentence Copy Typing | 80 |
| 93 | Closed Sentence Copy Typing | 100 |
| 106 | Closed Sentence Copy Typing | 100 |
| 107 | Closed Sentence Copy Typing | 130 |
